# Age-related disparities in oscillatory dynamics within scene-selective regions during spatial navigation

**DOI:** 10.1101/2023.10.16.562507

**Authors:** Marion Durteste, Alexandre Delaux, Ainhoa Ariztégui, Benoit R. Cottereau, Denis Sheynikhovich, Stephen Ramanoël, Angelo Arleo

## Abstract

Position is a key property that allows certain objects in the environment to attain navigational relevance. Symmetrical processing of object position across the horizontal meridian remains an unchallenged assumption of the spatial navigation literature. Nonetheless, a growing body of research reports vertical inhomogeneities in perceptual tasks, and recent evidence points towards a lower visual field preference in healthy aging. Factoring in the vertical position of objects to better apprehend spatial navigation abilities across the lifespan is essential. The occipital place area (OPA), parahippocampal place area (PPA), and medial place area (MPA) are ideal candidates to support the processing of the vertical location of navigational cues. Indeed, they are implicated in scene processing and spatial cognition, functions that may interact with their underlying retinotopic codes. This study aimed to elucidate whether young and older participants adjusted their gaze patterns and EEG activity within scene-selective regions in response to the vertical arrangement of landmarks. A sample of 21 young and 21 older participants completed a desktop-based task requiring them to navigate using objects at different vertical locations. We equipped them with an eye tracker and a high-density EEG cap. We used a fMRI-informed source-reconstruction algorithm to study the OPA, PPA and MPA with high spatial and temporal precision. Older adults exhibited a higher number of errors compared to young adults during reorientation. This age-related decline in accuracy was accompanied by a tendency among older participants to fixate objects in the lower half of the screen. This gaze bias was absent in young participants, who instead adapted their oculomotor behaviour based on the position of navigationally-relevant information. Another primary finding pertains to the differential pattern of theta and beta band activity in the OPA, PPA and RSC for environments that only contained relevant cues in the upper visual field. It thus appears that scene-selective regions process some aspect of the vertical position of information, irrespectively of their inherent retinotopic biases. Moreover, we revealed striking disparities between age groups in beta/gamma band synchronization in all scene-selective regions, indicating compromised top-down attentional mechanisms during spatial navigation in ageing. These age-related disparities in attentional dynamics might account for performance deficits in older participants. This study sheds light on a systematic downward gaze bias and altered attentional dynamics within scene-selective regions during spatial navigation in older age. It also emphasises the importance of considering vertical positioning as a fundamental property of objects within scenes.

## Introduction

Visual input is essential to forming accurate spatial representations that support successful wayfinding behaviour in humans (Ekstrom, 2015). In natural settings, objects often hold sufficient visuo-spatial information to serve as anchors for positional or directional knowledge thereby guiding future actions (Zhong and Moffat, 2016). One question that has sparked much debate in the field pertains to the properties which allow certain objects, among the abundance of incoming visual information, to attain navigational relevance (Chan *et al*., 2012). Intrinsic visual features that make objects useful for spatial navigation include size, colour, and distinctiveness (Stankiewicz and Kalia, 2007; Miller and Carlson, 2011; Auger, Mullally and Maguire, 2012; Auger and Maguire, 2018). Notwithstanding the contribution of these characteristics, non-physical properties such as location also play a decisive role in using a specific object as a spatial reference. A wealth of research has demonstrated that the position of objects in an environment conditions spatial behaviour. For example, the most recognizable and useful stimuli for navigation are those located at decision points (Janzen, 2006) or in close proximity to the goal (Waller and Lippa, 2007). Additionally, spatial representations differ according to the distance of objects with respect to the navigator. While proximal cues may be easier to rely upon as they engage action-based memories of learned trajectories, distal information enables the formation of precise cognitive maps (Hartley *et al*., 2003; Foo *et al*., 2005; Hurlebaus *et al*., 2008; Jabbari *et al*., 2021).

The undifferentiated processing of objects with respect to their location across the horizontal meridian nonetheless remains an unquestioned assumption in the literature (Hafed and Chen, 2016). We argue that position within the upper and lower visual fields represents an underappreciated cornerstone of visual information usage for spatial navigation. First, upper-lower inhomogeneities are pervasive across a wide array of perceptual and cognitive tasks. On the one hand, the lower visual field allows for better visual acuity, contrast sensitivity, spatial attention, and motion processing of stimuli (Regan, Erkelens and Collewijn, 1986; Skrandies, 1987; Carrasco, Talgar and Cameron, 2001; Rezec and Dobkins, 2004; Lakha and Humphreys, 2005; Levine and McAnany, 2005). The upper visual field, on the other hand, favors abilities such as visual search, change detection and categorical processing (Niebauer and Christman, 1998; Previc and Naegele, 2001; Rutkowski, Crewther and Crewther, 2002; Pflugshaupt *et al*., 2009). Second, the environment imposes differing constraints with respect to the type of information available for navigational purposes across the horizontal meridian. Monuments, immovable objects and wayfinding signage are typical components of the upper visual field whereas moving stimuli, obstacles, and spatial layout information appear predominantly in the lower visual field (Greene, 2013; Groen, Silson and Baker, 2017). In natural settings, one can therefore expect these vertical asymmetries to modulate both gaze dynamics and orienting of attention (Jóhannesson, Tagu and Kristjánsson, 2018). The above arguments provide solid grounds to factor in vertical position to the extensive literature on how object properties tie in with spatial behavior.

Wayfinding skills in the elderly are increasingly considered in light of the visual impairments that accompany healthy aging. Visuospatial processing declines in older age and modifies the way in which older adults sample the outside world and encode navigationally relevant visual information (Dowiasch *et al*., 2015; Zhong and Moffat, 2016; Kimura *et al*., 2019; Ramanoël *et al*., 2020). For example, a recent study found that healthy older adults’ orientation capabilities are rescued when geometry, instead of salient objects, is available to guide behavior (Bécu *et al*., 2023). Interestingly, accumulating evidence is pointing towards a possible influence of verticality on visuospatial function in healthy older adults. Vertical anisotropies in visual search appear to undergo significant changes throughout adulthood, shifting progressively to a lower visual field dominance (Brennan *et al*., 2017; Feng *et al*., 2017). Moreover, older adults have been found to gaze preferentially at the ground when reorienting in ecological laboratory settings (Bécu *et al*., 2020) and to show impaired spatial memory specifically for upper visual field objects (Durteste *et al*., 2023). Such reshaping of the use of visual space in healthy aging could have drastic consequences on the navigational relevance of objects located in the upper half of scenes.

At the neural level, the processing of scene layout and visual information contained within an environment has mainly been ascribed to the ventromedial posterior network of scene-selective regions. The occipital place area (OPA), parahippocampal place area (PPA) and medial place area (MPA) endorse distinct functions in this three-scene cortical system (Dilks, Kamps and Persichetti, 2022). The OPA underpins the processing of obstacles, available paths and egocentric motion thereby facilitating immediate visually guided navigation (Ferrara and Park, 2016; Kamps, Lall and Dilks, 2016; Bonner and Epstein, 2017). The PPA, by exploiting spatial layout as well as objects and their respective associations, is mostly responsible for the rapid categorization of a scene (Harel *et al*., 2016; Bilalić, Lindig and Turella, 2019). Finally, neural activity in the MPA (or retrosplenial cortex; see Silson, Steel and Baker, 2016) enables the bridging of local and global reference frames to support more complex map-based navigation (Marchette *et al*., 2014; Auger, Zeidman and Maguire, 2015; Persichetti and Dilks, 2019). The established roles of scene-selective regions in scene viewing and spatial cognition have been posited to interact with underlying retinotopic codes (Uyar *et al*., 2016; Steel *et al*., 2023). Indeed, the OPA, PPA and MPA inherit some of the retinotopic organization from early visual cortex and they exhibit preferences for designated areas of the visual field. The OPA and PPA display striking vertical biases for the lower visual field and upper visual field, respectively, while the MPA harbors a more general contralateral bias (Silson *et al*., 2015; Silson, Steel and Baker, 2016). The above findings make the OPA and PPA ideal candidates to support the encoding of the vertical position of objects during navigation. Research examining scene-selective regions in healthy aging is scarce but new evidence is pointing at a preserved connectivity and an increased activity of the OPA (Ramanoël *et al*., 2019, 2020). This result stands in contrast with the age-related alterations reported in the PPA and MPA (Ramanoël *et al*., 2015; Zhong and Moffat, 2018). No study has yet investigated whether such unequal aging of the cortical scene systems could alter the processing of information within the upper and lower visual fields of older adults in the context of navigation.

In the present study, we first test the hypothesis that the vertical position of objects modulates spatial navigation performance and does so differentially in young and older adult populations. Second, we sought to establish whether scene-selective regions in both age groups exhibit preferences for specific object positions, in line with their retinotopic biases. Most findings relating to the properties of scene-selective regions stem from experiments relying on the brief presentation of static scene images (Epstein and Baker, 2019), thus hindering the possibility to study top-down influences on their activity. We deemed it important that our paradigm allowed for unrestricted visual exploration and for scene processing in a navigational context. In this regard, we designed a desktop-based virtual navigation task coupled to an eye tracker that explored the behavioral and oculomotor patterns during orientation with cues of varying vertical location. To record cerebral activity pertaining to scene-selective regions, we used source-localized electroencephalography (EEG). Recent EEG works successfully extracted the timing at which scene-selective regions became sensitive to scene features underlining the added value of this technique in elucidating the underlying temporal dynamics (Kaiser, Turini and Cichy, 2019; Kaiser, Häberle and Cichy, 2020; Harel *et al*., 2022). To optimize the signal recovered from the OPA, PPA and MPA in participants, we opted for a region-of-interest (ROI) analysis using an individualized source reconstruction model informed with functional magnetic resonance imaging (fMRI) (Cottereau, Ales and Norcia, 2015).

## Results

To elucidate the role of the vertical position of information for spatial navigation, we asked 42 participants (21 young and 21 older adults) to perform a desktop-based virtual navigation task as we tracked their gaze patterns and EEG signals. Participants’ task was to learn the position of a goal in a square-like environment using the objects situated on the balconies and sidewalks at each intersection (Fig. 1). During the encoding phase, they watched videos of three trajectories to the goal. During the test phase, they viewed footage of multiple routes in the same environment and pressed a key upon arriving at each intersection to indicate the direction to the goal. We varied the position of navigationally relevant information across three different conditions. In the *UP* condition, only objects situated on balconies could be used to orient, while in the *DOWN* condition, only objects situated on sidewalks could be used. In the *MIX* condition, on the other hand, both balcony and sidewalk items were relevant for navigation. The experiment comprised 3 blocks, each divided into *UP*, *DOWN*, and *MIX* conditions and concluded with an object recognition task. To test our hypotheses, we compared young and older subjects’ behavioral performance, eye movement parameters and EEG activity across the three conditions of encoding and test phases of the navigational task. We used gaussian mixture modeling on participants’ fixation data to precisely delineate the upper and lower areas of interest (AOIs) and to conduct statistics on these regions (Fig. 1). We investigated dynamic cortical correlates of scene observation in the three ROIs (OPA, PPA, MPA) using Event-Related Spectral Perturbations (ERSPs) analysis on source-reconstructed EEG data. We looked for significant patterns in the ERSPs and their variations across age groups and conditions within each ROI. Guided by the main contrasts of the latter results, we conducted follow-up analyses to delineate ROI differences per age group within specific frequency bands.

**Figure 1.**
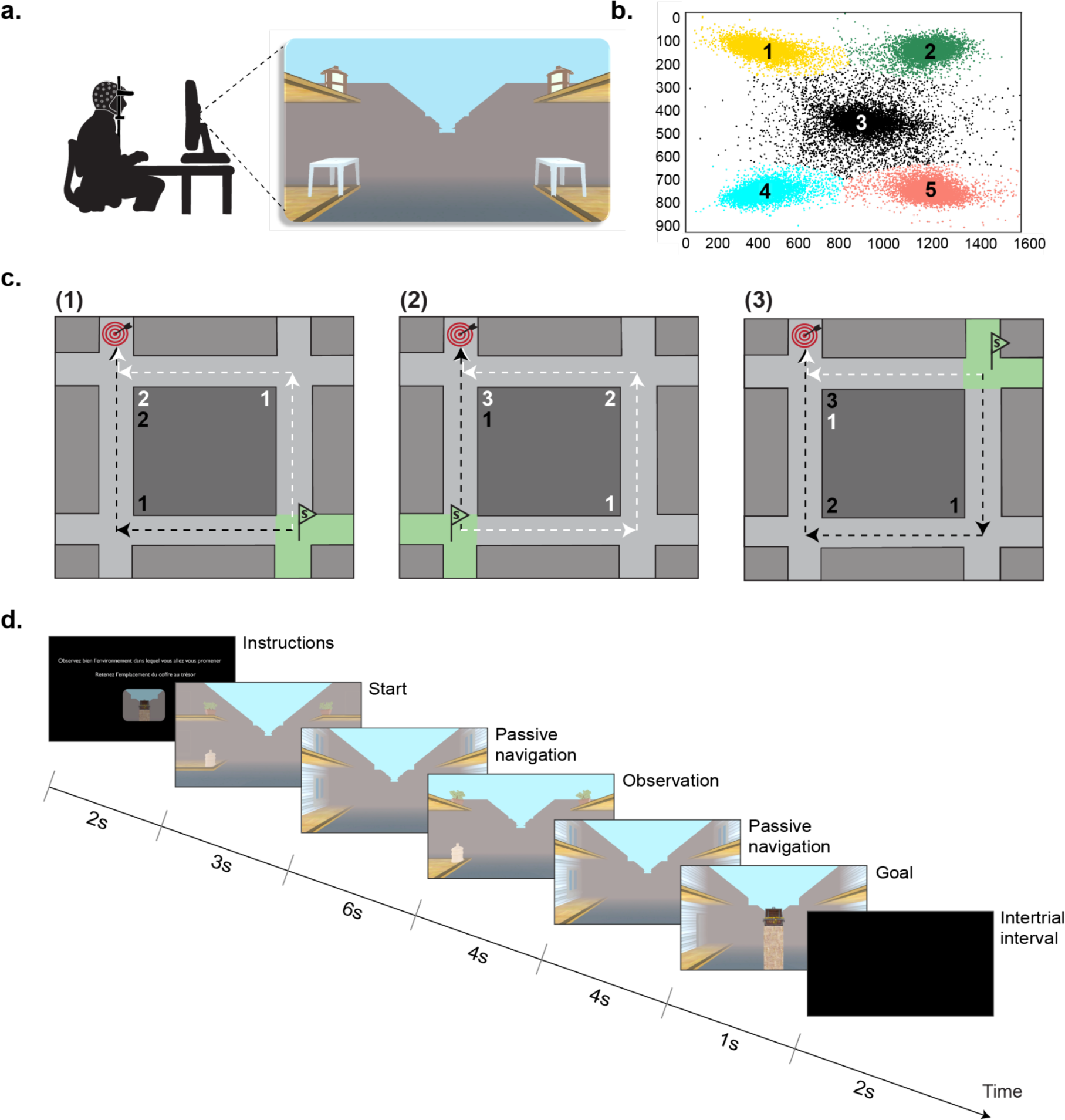
**a.** The participant, equipped with a 127-electrode EEG cap, is seated in front of a monitor with his head positioned on a head-mounted eye tracker. The screen displays a navigational task in a square-like virtual environment. **b.** Results from the GMM clustering analysis of fixation data at intersections during the encoding phase. All data points are depicted. The GMM defined five AOIs: top left (1), top right (2), center (3), bottom left (4), and bottom right (5). For data analysis, AOIs 1 and 2 are merged (upper AOI) and AOIs 4 and 5 are merged (lower AOI). Screen coordinates are in pixels. **c.** Schematic top view of the square-like layout of the virtual environment. The target represents a goal position, in the continuity of the top left intersection, within a specific instance of the environment. The green flags and green colored intersections mark the 3 starting positions associated with that goal. Each starting position is associated with two possible trajectories to the goal, that correspond to two distinct starting orientations. (1) If the participant starts from the bottom right intersection, it can take two routes that each pass through two intersections. (2, 3) If the participant starts from the bottom left or top left intersection, it can take two routes, one that passes through a single intersection and one that passes through three intersections. **d.** Schematic representation of a single encoding trial. Subjects navigate passively throughout the virtual environment and pause for 4 seconds at each intersection to encode the objects. The trial ends upon reaching the goal.

### Navigational behaviour varies as a function of age but not condition

We assessed participants’ performance on the spatial navigation task by computing the number of errors made during the test phase and by identifying the strategy used to reorient (mental map users vs. non mental map users). Across all three conditions, the average performance of young adults (M = 0.22 errors, SD = 0.22; t(62) = -28.67, *p* < 0.001, d = 3.16) and the average performance of older adults (M = 0.74 errors, SD = 0.19; t(62) = -11.27, *p* < 0.001, d = 1.42) were significantly above the chance level of 1 error per intersection. A chi-square test of independence revealed that there was a significant association between age and strategy (*χ*^2^ (1, *N* = 42) = 16.43, *p* < 0.001). A significantly greater proportion of young participants (19/21) reported being mental map users compared with older participants (5/21). A second chi-square test showed a significant relationship between age and object selection (*χ*^2^(1, *N* = 42) = 11.76, *p* < 0.001). Older adults reported relying on the 4 objects at intersections (15/21) in significantly greater proportion than young participants (3/21) who were more inclined to select a subset of objects. We then ran a linear mixed model aimed at explaining the number of errors with participants as random intercepts and age, sex, condition, and strategy as fixed effects. Interactions between age, condition and strategy were discarded as they resulted in worse model fits. First, we found that age had a significant influence on navigational performance (F(1, 37) = 17.98, *p* < 0.001, *η_p_^2^* = 0.33, 95 % Confidence Interval (CI) = [0.10, 0.53]; Fig. 2b) with older subjects making more errors than young participants per intersection (older: M = 0.74, SD = 0.19; young: M = 0.22, SD = 0.22). Moreover, there was a main effect of sex on the number of mistakes made during the test phase (F(1, 37) = 12.95, *p* < 0.001, *η_p_^2^* = 0.26, 95 % CI = [0.054, 0.47]) with a higher mean number per intersection reported in females (M = 0.60, SD = 0.27) than in males (M = 0.39, SD = 0.33). We also found an influence of strategy on navigational performance (F(1, 37) = 10.97, *p* = 0.0021, *η_p_^2^* = 0.23, 95 % CI = [0.037, 0.44]). Across age groups, individuals who did not use a mental map to reorient (M = 0.74, SD = 0.18) displayed significantly worse performance than those who did (M = 0.28, SD = 0.27). The linear mixed model did not reveal any evidence for an effect of condition on navigation performance (F(2, 208) = 0.50, *p* = 0.61, *η_p_^2^* = 0.0048, 95 % CI = [0.00, 0.033]). We ran a second linear mixed model with participants as random intercepts to investigate the impact of repeated *vs.* new test trials on behavioral performance. Our results indicated that there was a significant interaction between age and type of test trial on performance (F(1, 208) = 8.38, *p* = 0.0042, *η_p_^2^* = 0.039, 95 % CI = [0.0040, 0.10]; Fig. 2c). We found a significant difference between repeated and new routes in older adults only (t(208) = 1.57, *p* = 0.0096, d = 0.37, SE = 0.18), with new routes leading to a greater average number of mistakes (M = 0.77, SD = 0.16) compared with repeated routes (M = 0.70, SD = 0.18).

**Figure 2.**
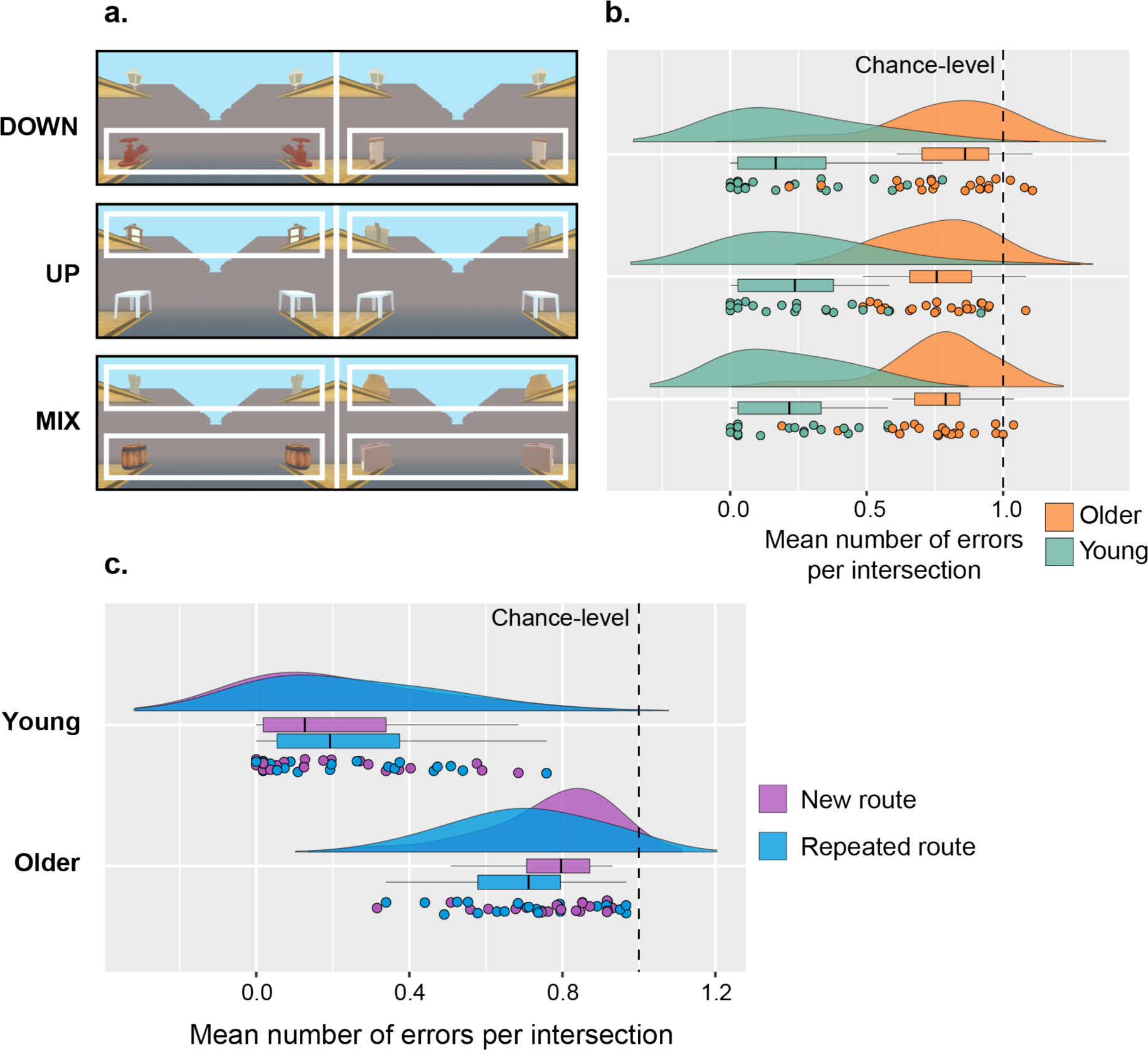
**a.** Screenshots of two intersections of a DOWN, UP, and MIX condition. The white boxes highlight the objects that are navigationally relevant in each condition. Note that in a MIX condition all objects are helpful to orient. **b**. Distributions of the mean number of errors per intersection according to the condition in young and older adults. Age has a significant influence on navigational performance (F(1, 37) = 28.57, *p* < 0.001, *η_p_^2^* = 0.44, 95 % Confidence Interval (CI) = [0.20, 0.61]). **c.** Distributions of the mean number of errors per intersection for a new or repeated route in young and older adults. Older adults perform better on repeated routes than on new routes (t(208) = 1.57, *p* = 0.0096, d = 0.37, SE = 0.18).

Finally, we assessed associations between performance on the spatial navigation task and neuropsychological test scores. The overall linear regression models for the perspective-taking (R^2^ = 0.65, F(2, 39) = 35.61, *p* < 0.001) and 3D mental rotation tasks (R^2^ = 0.64, F(2, 36) = 31.76, *p* < 0.001) were statistically significant. We found that lower performance on the perspective-taking (β = 0.073, *p* = 0.011, *η_p_^2^* = 0.15, 95 % CI = [0.0087, 0.36]) and on the 3D mental rotation (β = -0.37, *p* = 0.022, *η_p_^2^* = 0.14, 95 % CI = [0.0014, 0.35]) tasks were associated with a significantly greater number of navigational errors across all participants.

### Adapting gaze patterns to the condition is impaired in older age

#### Gaze patterns during the encoding phase

We first examined participants’ gaze patterns across all 4-second observation periods of the encoding phase. We ran separate linear mixed models to study the effects of age, condition, and sex on dwell time in the lower, upper, and central AOI. Regarding the proportion of time spent looking at the lower AOI, we found little evidence for main effects of age (F(1, 38) = 1.25, *p* = 0.27, *η_p_^2^* = 0.032, 95 % CI = [0.00, 0.20]) or sex (F(1, 38) = 3.67, *p* = 0.063, *η_p_^2^* = 0.088, 95 % CI = [0.00, 0.29]). However, we uncovered a significant influence of condition on dwell time in the lower AOI (F(2, 317) = 3.60, *p* = 0.020, *η_p_^2^* = 0.022, 95 % CI = [0.00, 0.060]). Post-hoc comparisons showed a significant difference between the DOWN and MIX conditions (t(317) = -2.90, *p* = 0.035, d = 0.24, SE = 0.13), with longer time spent looking at the lower AOI in the MIX condition (M = 37.02%, SD = 11.73%) than in the DOWN condition (M = 34.12%, SD = 12.77%) across all participants. We then examined the upper AOI and we found no evidence that age (F(1, 38) = 0.83, *p* = 0.37, *η_p_^2^* = 0.021, 95 % CI = [0.00, 0.17]) or sex (F(1, 38) = 0.74, *p* = 0.40, *η_p_^2^* = 0.010, 95 % CI = [0.00, 0.15]) significantly predicted dwell time in that area. There was a significant main effect of condition on dwell time in the upper AOI (F(2, 315) = 6.03, *p* = 0.0027, *η_p_^2^* = 0.037, 95 % CI = [0.0049, 0.083]). Post-hoc Tukey testing revealed that upper dwell time was significantly different between the MIX and UP conditions (t(315) = 3.70, *p* = 0.0040, d = 0.31, SE = 0.13) and between the MIX and DOWN conditions (t(315) = -3.13, *p* = 0.018, d = 0.24, SE = 0.13). Participants spent longer fixating the upper AOI during the MIX condition (M = 35.34%, SD = 13.46%) than during the UP (M = 31.30%, SD = 12.80%) and DOWN conditions (M = 32.11%, SD = 13.73%). Finally, there was little evidence to support a main effect of age (F(1, 38) = 0.0004, *p* = 0.98, *η_p_^2^* = 0.00, 95 % CI = [0.00, 0.00]) on central dwell time. We nonetheless found that sex (F(1, 38) = 4.55, *p* = 0.040, *η_p_^2^* = 0.11, 95 % CI = [0.00, 0.31]) and condition (F(2, 315) = 13.21, p < 0.001, *η_p_^2^* = 0.077, 95 % CI = [0.028, 0.14]) had significant influences on time spent looking at the central AOI. We uncovered that central dwell time in male participants (M = 35.65%, SD = 14.81%) was superior to central dwell time in female participants (M = 27.15%, SD = 16.90%). Post-hoc comparisons also showed significant differences in central dwell time between the MIX and UP conditions (t(315) = - 6.17, *p* < 0.001, d = 0.41, SE = 0.13) and between the MIX and DOWN conditions (t(315) = 6.16, *p* < 0.001, d = 0.40, SE = 0.13), with all participants fixating the central AOI longer during the UP (M = 34.14%, SD = 17.50%) and DOWN conditions (M = 33.77%, SD = 16.23%) than during the MIX condition (M = 27.64%, SD = 14.38%). We ran another linear mixed model to investigate the influence of age, sex, and condition on the VMA for dwell time. The VMA computes the vertical asymmetry in the distribution of time spent looking at the AOIs. We found no evidence for main effects of age (F(1, 38) = 2.02, *p* = 0.16, *η_p_^2^* = 0.050, 95 % CI = [0.00, 0.23]), sex (F(1, 38) = 0.34, *p* = 0.57, *η_p_^2^* = 0.012, 95 % CI = [0.00, 0.19]), or condition (F(2, 317) = 0.19, p = 0.83, *η_p_^2^* = 0.0012, 95 % CI = [0.00, 0.013]) on the VMA for dwell time. The distribution of time spent looking across the upper and lower AOIs remains stable across conditions. One-sample Wilcoxon signed rank tests showed that the mean VMA per condition per block was significantly different from zero in older participants (V(182) = 11387, *p* < 0.001, median = 18.52, 95% CI [10.07, 26.76], r = 0.31) but not in young participants (V(178) = 7768, *p* = 0.68, median = -2.00, 95% CI [-11.31, 6.85], r = 0.031).

Finally, we ran linear mixed models to study the proportion of first fixations across all intersections of the encoding phase directed towards the lower, upper, or central AOIs. We stress that the term *first fixation* refers to the very first fixation upon arrival at each intersection, *i.e.,* the first fixated AOI at each intersection. We found no evidence to support a significant influence of age (F(1, 39) = 0.0002, *p* = 0.99, *η_p_^2^* = 0.00, 95 % CI = [0.00, 0.00]), sex (F(1, 39) = 1.66, *p* = 0.20, *η_p_^2^* = 0.041, 95 % CI = [0.00, 0.21]) or condition (F(2, 325) = 1.10, *p* = 0.33, *η_p_^2^* = 0.0068, 95 % CI = [0.00, 0.031]) on the proportion of first fixations directed towards the lower AOI. Moreover there was no evidence for statistically significant main effects of sex (F(1, 39) = 0.63, *p* = 0.43, *η_p_^2^* = 0.016, 95 % CI = [0.00, 0.16]) or condition (F(2, 323) = 1.42, *p* = 0.24, *η_p_^2^* = 0.0087, 95 % CI = [0.00, 0.035]) on the proportion of first fixations located in the upper AOI. We found that age (F(1, 39) = 5.22, *p* = 0.028, *η_p_^2^* = 0.12, 95 % CI = [0.00, 0.32]) and the interaction between age and condition (F(2, 323) = 3.56, *p* = 0.030, *η_p_^2^* = 0.022, 95 % CI = [0.00, 0.059]; Figs. 3a,b) significantly predicted the proportion of upper first fixations. Young participants had a greater proportion of first fixations directed towards the upper AOI than did older participants across conditions (young: M = 34.57%, SD = 23.84%; older: M = 21.93%, SD = 22.14%). Furthermore, post-hoc tests revealed an absence of significant pairwise comparisons for the interaction of age and condition. There was no evidence to indicate that sex (F(1, 39) = 0.28, *p* = 0.60, *η_p_^2^* = 0.0072, 95 % CI = [0.00, 0.14]) significantly predicted the proportion of first fixations situated in the central AOI. However, we did reveal main effects of age (F(1, 39) = 7.65, *p* = 0.0086, *η_p_^2^* = 0.16, 95 % CI = [0.012, 0.37]) and condition (F(2, 323) = 3.72, *p* = 0.025, *η_p_^2^* = 0.023, 95 % CI = [0.00, 0.060]). Older participants looked at the central cluster at the start of each intersection to a significantly greater extent than young participants (young: M = 39.18%, SD = 18.69%; older: M = 53.43%, SD = 22.76%). We also found the interaction between age and condition to be significant (F(2, 323) = 3.71, *p* = 0.022, *η_p_^2^* = 0.022, 95 % CI = [0.00, 0.060]). Post-hoc Tukey tests showed a significant age-related difference in the proportion of central first fixations during the UP condition only (t(55) = 17.47, *p* = 0.023, d = 0.89, SE = 0.19). Indeed, older participants fixated the central AOI at the start of UP intersections to a greater extent than did young participants (young: M = 38.24%, SD = 16.57%; older: M = 56.11%, SD = 23.18%). We also found a significant difference between the DOWN and MIX conditions in young adults (t(323) = 8.60, *p* = 0.012, d = 0.45, SE = 0.18). Young participants had a significantly greater proportion of first fixations located in the central AOI during the DOWN condition (M = 43.94%, SD = 21.62%) than during the MIX condition (M = 35.34%, SD = 16.64%).

**Figure 3.**
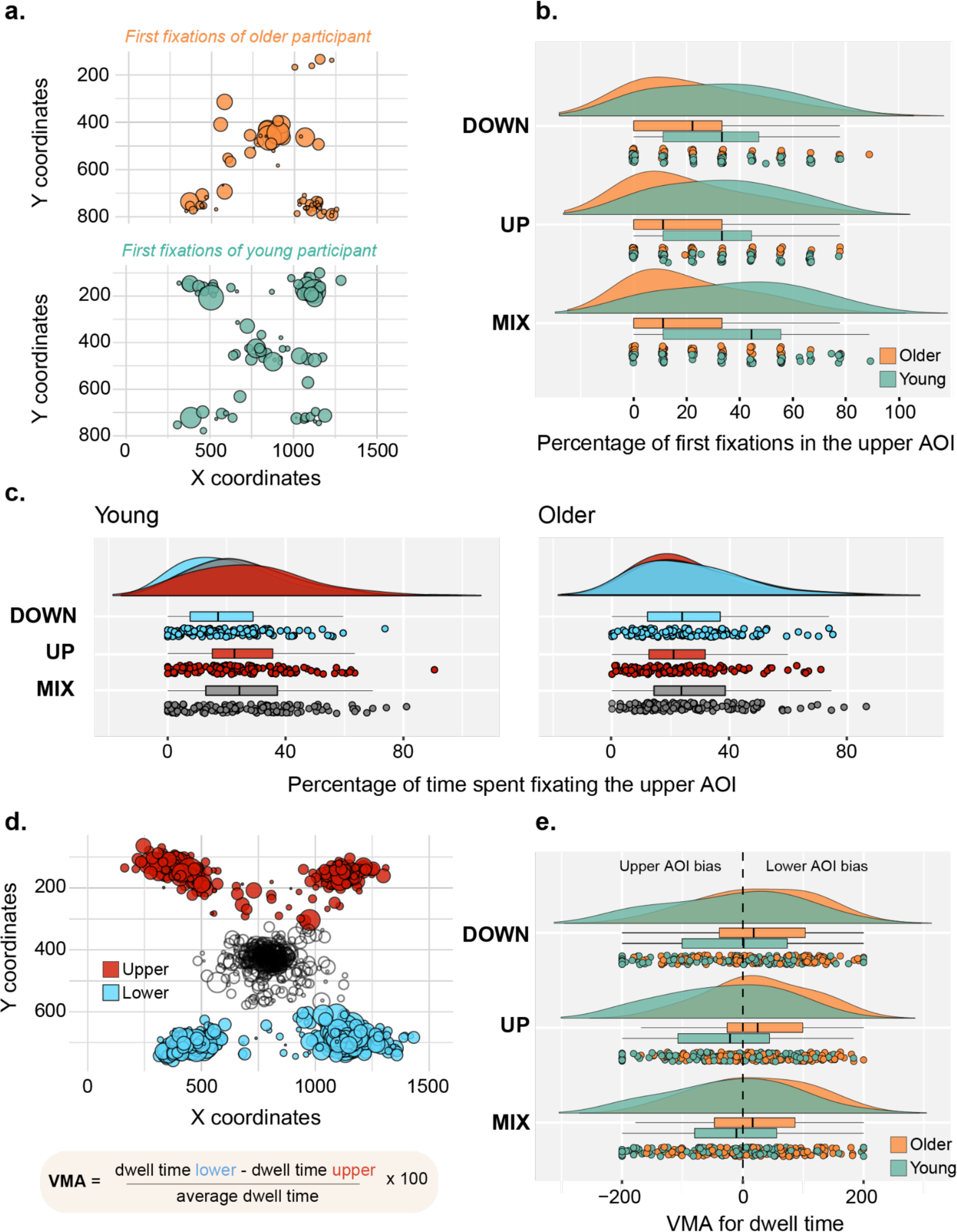
Eye movement parameters in young and older participants. **a**. Depiction of all first fixations at the start of intersections during the encoding phase for an example older participant and an example young participant. Dots represent fixations and their size covaries with the duration of the fixation. **b**. Distributions of the percentage of first fixations situated in the upper area-of-interest (AOI) in young and older adults across conditions during the encoding phase. Age is a significant predictor of first fixations in the upper AOI (F(1, 39) = 5.22, *p* = 0.028, *η_p_^2^* = 0.12, 95 % CI = [0.00, 0.32]). **c**. Distributions of the percentage of time spent fixating the upper AOI in young and older adults across conditions during the 4 seconds after arriving at the intersection in the test phase. There were significant differences in upper dwell time between the DOWN and MIX conditions (t(677) = -7.76, *p* < 0.001, d = 0.47, SE = 0.13) and between the DOWN and UP conditions (t(677) = -5.75, *p* < 0.001, d = 0.35, SE = 0.13) in young adults only. **d**. Depiction of all fixations during the 4 seconds after arriving at the intersection in the test phase for an example participant. The vertical meridian asymmetry (VMA) computes the difference in dwell time between the lower and upper AOIs. Indices that are close to 0 indicate no asymmetry in dwell time between the upper and lower AOIs, indices that are superior to 0 indicate a bias for the lower AOI, and indices that are inferior to 0 indicate a bias for the upper AOI. Dots represent fixations and their size covaries with the duration of the fixation. **e**. Distributions of the VMA for dwell time in young and older adults across conditions during the test phase. Age is a significant predictor of the VMA (F(1, 38) = 4.70, *p* = 0.036, *η_p_^2^* = 0.11, 95 % CI = [0.00, 0.31]).

#### Gaze patterns during the test phase

We then focused our analysis on participants’ gaze patterns across all pre-decision intervals of the test phase. The latter correspond to the periods between arrival at the intersection and pressing a key to indicate the correct direction of travel. They were restricted to a maximum of 4 seconds even if participants took longer to make a decision. We ran separate linear mixed models to test the influence of age, condition, and sex on the proportion of time spent looking at the upper, lower, and central AOIs. We found no evidence to support a main effect of age (F(1, 38) = 0.072, *p* = 0.79, *η_p_^2^* = 0.0019, 95 % CI = [0.00, 0.10]) on the proportion of upper dwell time. There were, however, statistically significant influences of sex (F(1, 38) = 4.69, *p* = 0.037, *η_p_^2^* = 0.11, 95 % CI = [0.00, 0.31]), condition (F(2, 677) = 10.76, *p* < 0.001, *η_p_^2^* = 0.031, 95 % CI = [0.0093, 0.059]), and the interaction between age and condition (F(2, 677) = 8.56, *p* < 0.001, *η_p_^2^* = 0.025, 95 % CI = [0.0058, 0.051]; Fig. 3c) on the proportion of time spent fixating the upper AOI. Indeed, dwell time in the upper AOI was superior in female participants (M = 30.05%, SD = 17.84%) than in male participants (M = 21.37%, SD = 14.65%). Post-hoc Tukey tests showed that there were significant differences in upper dwell time between conditions in young adults only, between the DOWN and MIX conditions (t(677) = -7.76, p < 0.001, d = 0.47, SE = 0.13) and between the DOWN and UP conditions (t(677) = -5.75, p < 0.001, d = 0.35, SE = 0.13). Young participants spent less time fixating the upper AOI during the DOWN condition (M = 20.58%, SD = 15.03%) than during the MIX (M = 28.34%, SD = 17.84%) and UP conditions (M = 26.12%, SD = 16.61%). Examining lower dwell time, we found no evidence for significant main effects of age (F(1, 38) = 4.09, *p* = 0.050, *η_p_^2^* = 0.097, 95 % CI = [0.00, 0.30]) or sex (F(1, 38) = 1.76, *p* = 0.19, *η_p_^2^* = 0.044, 95 % CI = [0.00, 0.22]). We nonetheless revealed that condition was a significant predictor of the proportion of time spent in the lower AOI (F(2, 679) = 3.22, *p* = 0.041, *η_p_^2^* = 0.0094, 95 % CI = [0.00, 0.027]). There were significant differences in lower dwell time between the DOWN and MIX conditions (t(679) = -2.49, *p* = 0.031, d = 0.14, SE = 0.091) across all participants. They spent more time looking at the lower AOI during the MIX condition (M = 28.13%, SD = 17.45%) than during the DOWN condition (M = 25.63%, SD = 17.84%). Moreover, the linear mixed model analyzing dwell time in the central AOI revealed no evidence that age was a significant predictor (F(1, 38) = 1.99, *p* = 0.17, *η_p_^2^* = 0.050, 95 % CI = [0.00, 0.23]). However, we found main effects of sex (F(1, 38) = 7.06, *p* = 0.011, *η_p_^2^* = 0.16, 95 % CI = [0.0086, 0.37]), condition (F(2, 677) = 17.67, *p* < 0.001, *η_p_^2^* = 0.050, 95 % CI = [0.021, 0.084]), and the interaction between age and condition (F(2, 677) = 10.30, *p* < 0.001, *η_p_^2^* = 0.030, 95 % CI = [0.0085, 0.057]) on central dwell time. Male participants spent significantly more time fixating the central AOI than did female participants (male: M = 55.21%, SD = 20.96%; female: M = 38.91%, SD = 21.45%). In addition, post-hoc pairwise comparisons showed central dwell time differences between the DOWN and MIX conditions (t(677) = 11.98, *p* < 0.001, d = -0.52, SE = 0.13) and between the DOWN and UP conditions (t(677) = 7.49, *p* < 0.001, d = -0.31, SE = 0.13) in young adults only. Indeed, young participants spent significantly more time looking at the central AOI during the DOWN condition (M = 59.34%, SD = 23.77%) than during the MIX (M = 47.35%, SD = 22.33%) and UP conditions (M = 52.15%, SD = 23.25%). To complement the analyses regarding dwell time proportions, we ran a linear mixed model examining the influence of age, sex, and condition on the VMA for dwell time. Whereas we found no evidence to suggest main effects of sex (F(1, 38) = 0.43, *p* = 0.51, *η_p_^2^* = 0.011, 95 % CI = [0.00, 0.15]) or condition (F(2, 670) = 0.37, *p* = 0.69, *η_p_^2^* = 0.0011, 95 % CI = [0.00, 0.0089]) on the VMA, we did show that age (F(1, 38) = 4.70, *p* = 0.036, *η_p_^2^* = 0.11, 95 % CI = [0.00, 0.31]; Figs 3d,e) was a significant predictor of the VMA. The VMA in older adults (M = 23.74, SD = 88.20) was significantly higher than the VMA in young adults (M = -20.35, SD = 101.72), underlining a greater fixation bias towards the lower AOI in older age. One-sample Wilcoxon signed rank tests revealed that the mean VMA per condition per block was significantly different from zero in young (V(178) = 5736, *p* = 0.023, median = -17.67, 95% CI [-32.34, -2.42], r = 0.17) and in older participants (V(183) = 11029, *p* < 0.001, median = 24.76, 95% CI [11.65, 37.40], r = 0.27).

We ran additional linear mixed models to study the proportion of first fixations across all intersections of the test phase situated in the lower, upper, or central AOIs. We reported no statistical evidence for main effects of age (F(1, 39) = 1.09, *p* = 0.30, *η_p_^2^* = 0.027, 95 % CI = [0.00, 0.19]), sex (F(1, 39) = 0.48, *p* = 0.49, *η_p_^2^* = 0.012, 95 % CI = [0.00, 0.15]) or condition (F(2, 327) = 2.57, *p* = 0.078, *η_p_^2^* = 0.015, 95 % CI = [0.00, 0.048]) on the proportion of first fixations located in the lower AOI. Regarding first fixations towards the upper AOI, we found no evidence to support influences of age (F(1, 39) = 0.33, *p* = 0.57, *η_p_^2^* = 0.0083, 95 % CI = [0.00, 0.14]) or sex (F(1, 39) = 1.71, *p* = 0.20, *η_p_^2^* = 0.042, 95 % CI = [0.00, 0.22]). However, we did reveal that condition predicted the proportion of upper first fixations (F(2, 327) = 6.35, *p* = 0.0020, *η_p_^2^* = 0.037, 95 % CI = [0.0056, 0.083]). Post-hoc Tukey testing revealed significant differences between the DOWN and UP conditions (t(325) = -5.16, p = 0.0026, d = 0.23, SE = 0.13) as well as between the DOWN and MIX conditions (t(325) = -4.18, p = 0.020, d = 0.20, SE = 0.13). Across age groups, participants directed less first fixations towards the upper AOI during the DOWN condition (M = 21.62, SD = 19.81) than during the UP (M = 26.56, SD = 23.46) and MIX conditions (M = 25.80, SD = 22.88). Finally, while we did not find any evidence for a main effect of age (F(1, 39) = 0.20, *p* = 0.66, *η_p_^2^* = 0.0051, 95 % CI = [0.00, 0.13]) or sex (F(1, 39) = 0.42, *p* = 0.52, *η_p_^2^* = 0.011, 95 % CI = [0.00, 0.15]), we uncovered a main effect of condition (F(2, 325) = 13.72, *p* < 0.001, *η_p_^2^* = 0.078, 95 % CI = [0.029, 0.14]), and a significant interaction between age and condition (F(2, 325) = 5.53, *p* = 0.0044, *η_p_^2^* = 0.033, 95 % CI = [0.0036, 0.076]) on the proportion of first fixations situated in the central AOI. Post-hoc Tukey testing showed differences between the DOWN and UP conditions (t(323) = 13.26, *p* < 0.001, d = 0.51, SE = 0.18) and between the DOWN and MIX conditions (t(323) = 10.89, *p* < 0.001, d = 0.43, SE = 0.18) in young adults only. At the start of each intersection, young participants had a greater proportion of fixations that were situated in the central AOI during the DOWN condition (M = 57.59%, SD = 26.75%) than during the UP (M = 44.87%, SD = 23.46%) and MIX conditions (M = 46.70%, SD = 24.17%).

### EEG activity in scene-selective regions differs across age groups and conditions

To enable the most accurate spatial allocation of EEG activity, we relied on a fMRI informed source reconstruction model tailored to each participant (Cottereau, Ales and Norcia, 2012, 2015). Individual data included MRI anatomical scans for head and brain structure, a fMRI localizer task for scene-selective regions (Ramanoël *et al*., 2020) and a 3D model of electrode positions co-registered with the anatomical MRI data. We solved the inverse problem using the L2-regularized minimum-norm estimation (MNE) model (Hämäläinen *et al*., 1993).

We conducted an ERSP analysis time-locked to the event of arriving at an intersection (t = 0 ms) and grouped by principal frequency bands (delta: <4 Hz, theta: 4-8 Hz, alpha: 8-12 Hz, beta: 12-30 Hz, gamma: >30 Hz). We averaged the source-reconstructed power spectrum from both hemispheres to summarize bilateral ROI activity, as we did not have any hypothesis on lateralized effects. We reported average ERSPs baselined with the activity preceding the arrival at the level of an intersection. This baseline corresponds to participants passively navigating through a street that is devoid of any relevant spatial information. We computed ERSP statistical differences with permutation tests based on the maximum cluster-level statistic (Maris and Oostenveld, 2007) using 1,000 permutations. We set the initial significance level to *p* < 0.05, and Bonferroni-corrected for variables with more than 2 levels.

#### Encoding trials

The investigated ERSP activity during the encoding trials relates to the perception of the intersection and the objects contained within it. We allowed participants a fixed 4-second static observation period at each intersection to give them enough time to scan the whole scene and encode the spatial relationships between intersections (see Fig. 1d). We found ERSP activity during the encoding trials to be highly similar to that associated with the test trials, particularly with regards to the patterns of average activity and age-related differences. Worthy of note, there are half as many trials in the encoding phase as in the test phase, which increases statistical power in the latter. We therefore chose to only present results relative to test trials and we refer the interested reader to the Supplementary Figs 1, 2 and 3 for a thorough description of ERSP activity during the encoding phase.

#### Test trials

The investigated ERSP activity during test trials relates to the observation phase at the intersection prior to the navigational decision. Therefore, this static observation and decision period depended on the time it took for participants to choose the correct direction. To be consistent with the fixed observation time during the encoding trials, we inspected the first 4 seconds of cortical activity following arrival at the intersection.

##### ERSP activity in the ROIs

First, we found a sustained synchronization in the delta/theta frequency band (2-8 Hz) across all ROIs in both age groups, starting 250 ms before event onset and sustained over the 4-second window (Figs. 4a, 5a, 6a). It was more pronounced during the first second of observation, particularly in the older adult group. Indeed, we reported a significant increase in synchronization in the theta band between +100 ms and +750 ms after arrival at the intersection in the OPA (Fig. 4c) and MPA of older participants (Fig. 6c).

**Figure 4.**
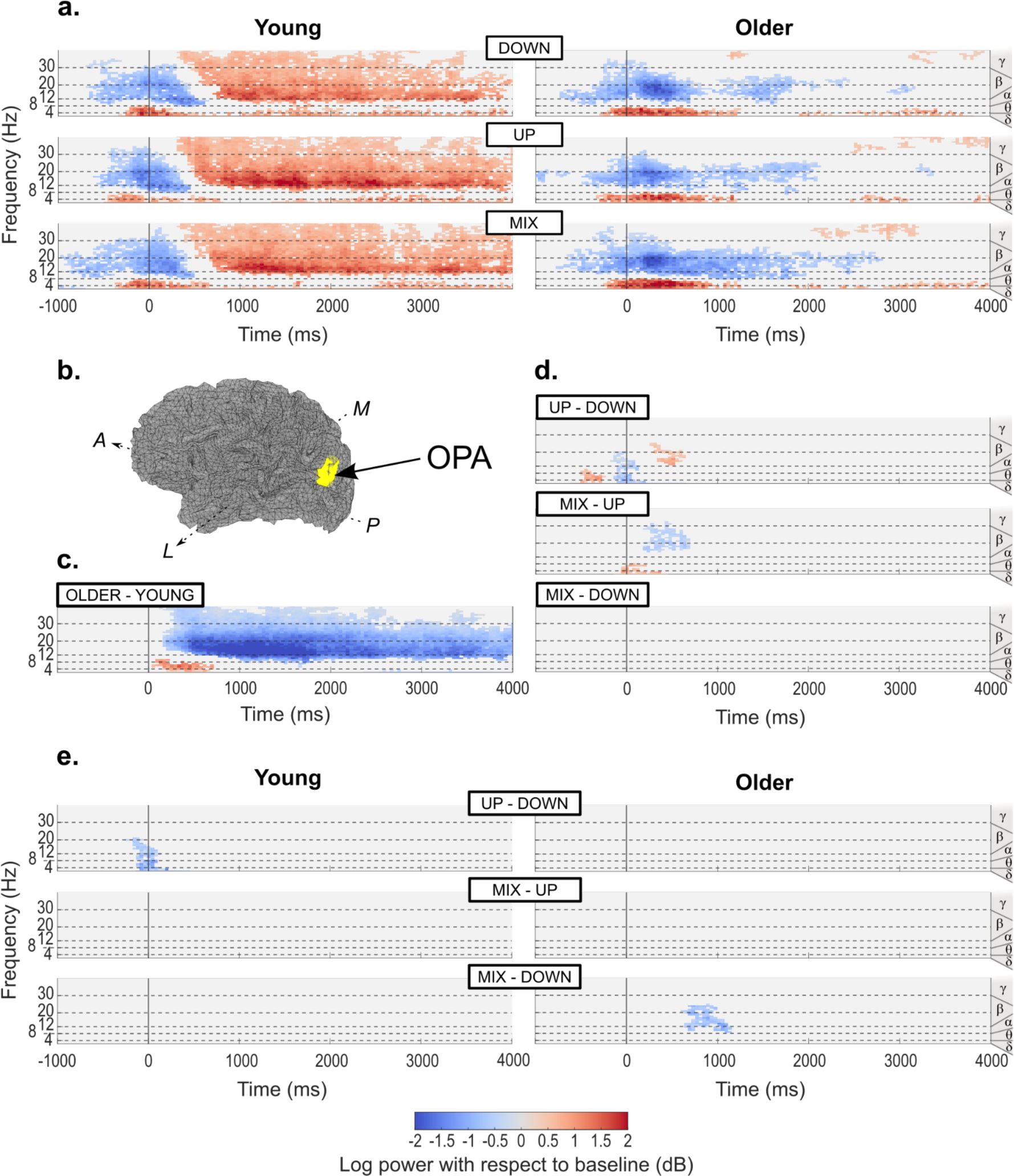
ERSP activity time-locked to the event of arriving at an intersection in the test trials, reconstructed in the OPA. Activity merged from both hemispheric locations after source reconstruction. We examined delta (δ; <4Hz), theta (θ; 4-8Hz), alpha (α; 8-12Hz), beta (β; 12-30Hz) and gamma (γ; >30Hz) frequency bands. We investigated all statistical differences with permutation tests based on the maximum cluster-level statistic using 1,000 permutations. We used linear mixed-effects modeling to evaluate the statistical significance at the “pixel” level (spectral power at a given time-frequency pair) for a given permutation. **a**. Average activity per age group and condition baselined with the 2 seconds period prior to arrival at the intersection. We only display activity statistically different from surrogate baseline distribution (*p* < 0.05). **b**. Illustration of the localization of the left OPA overlaid on left hemisphere source space (midgray surface) in one young participant. A: Anterior; P: Posterior; M: Medial; L: Lateral. **c**. Differences between age groups, irrespective of conditions. We indicate the direction of the difference above each graph (*e.g.* for the graph entitled “OLDER-YOUNG”, we subtracted the average signal for the young adults from the average signal for the older adults, such that a positive difference means more power in the older group than in the young group and a negative difference means less power in the older group than in the young group). We only display statistically significant differences (*p* < 0.05).**d**. Differences between conditions, irrespective of age groups. We only display statistically significant differences (*p* < 0.0083, Bonferroni-corrected for 3 two-sided comparisons). **e**. Differences between conditions within each age group. We only display statistically significant differences (*p* < 0.0041, Bonferroni-corrected for 6 two-sided comparisons).

**Figure 5.**
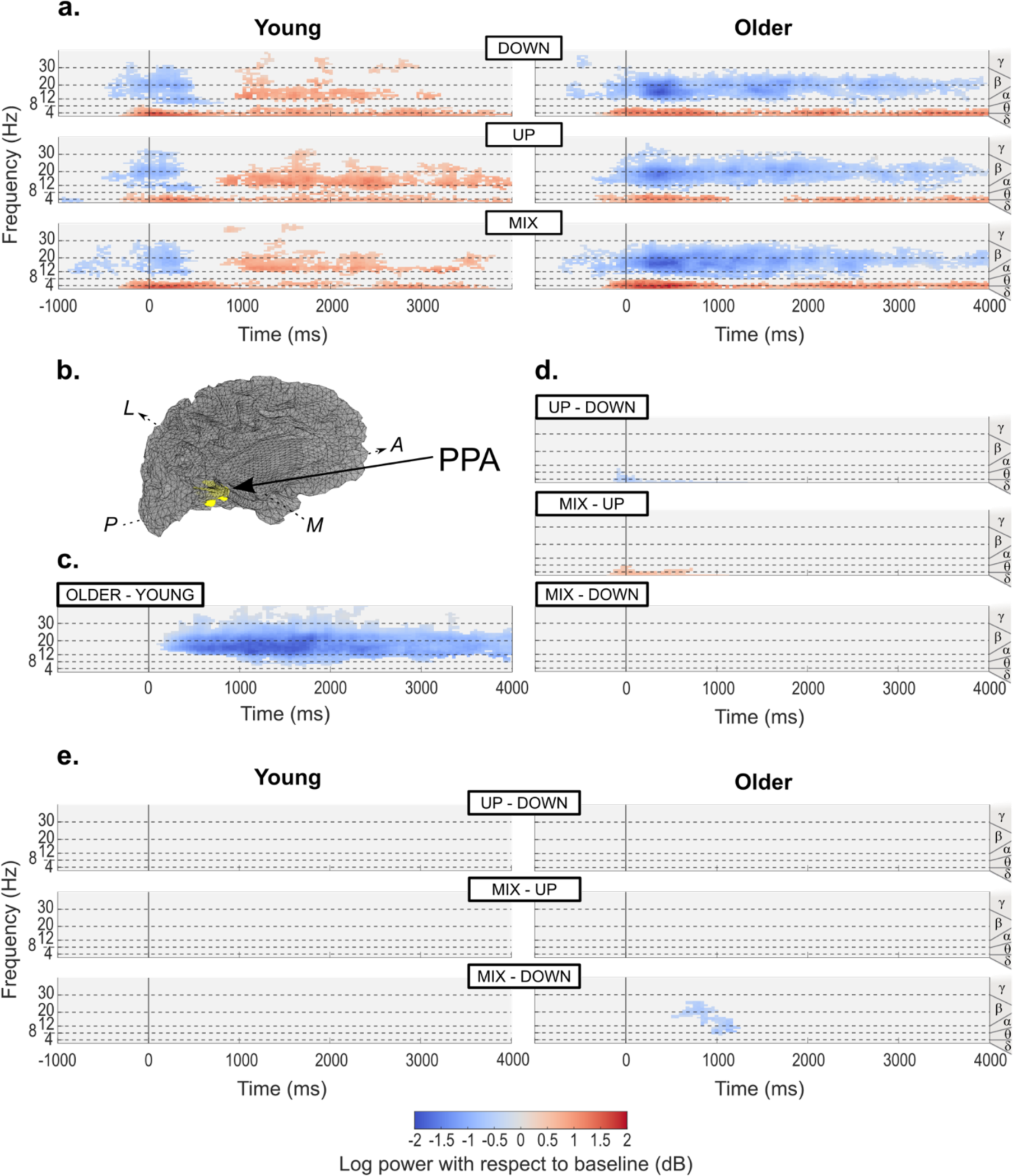
ERSP activity time-locked to the event of arriving at an intersection in the test trials, reconstructed in the PPA. For further details, see Fig. 4 legend. **a**. Average activity per age group and condition baselined with the 2 seconds period prior to arrival at the intersection. We only display activity statistically different from surrogate baseline distribution (*p* < 0.05). **b**. Illustration of the localization of the left PPA overlaid on left hemisphere source space (midgray surface) in one young participant. A: Anterior; P: Posterior; M: Medial; L: Lateral. **c**. Differences between age groups, irrespective of conditions. We only display statistically significant differences (*p* < 0.05). **d**. Differences between conditions, irrespective of age groups. We only display statistically significant differences (*p* < 0.0083, Bonferroni-corrected for 3 two-sided comparisons). **e**. Differences between conditions within each age group. We only display statistically significant differences (*p* < 0.0041, Bonferroni-corrected for 6 two-sided comparisons).

**Figure 6.**
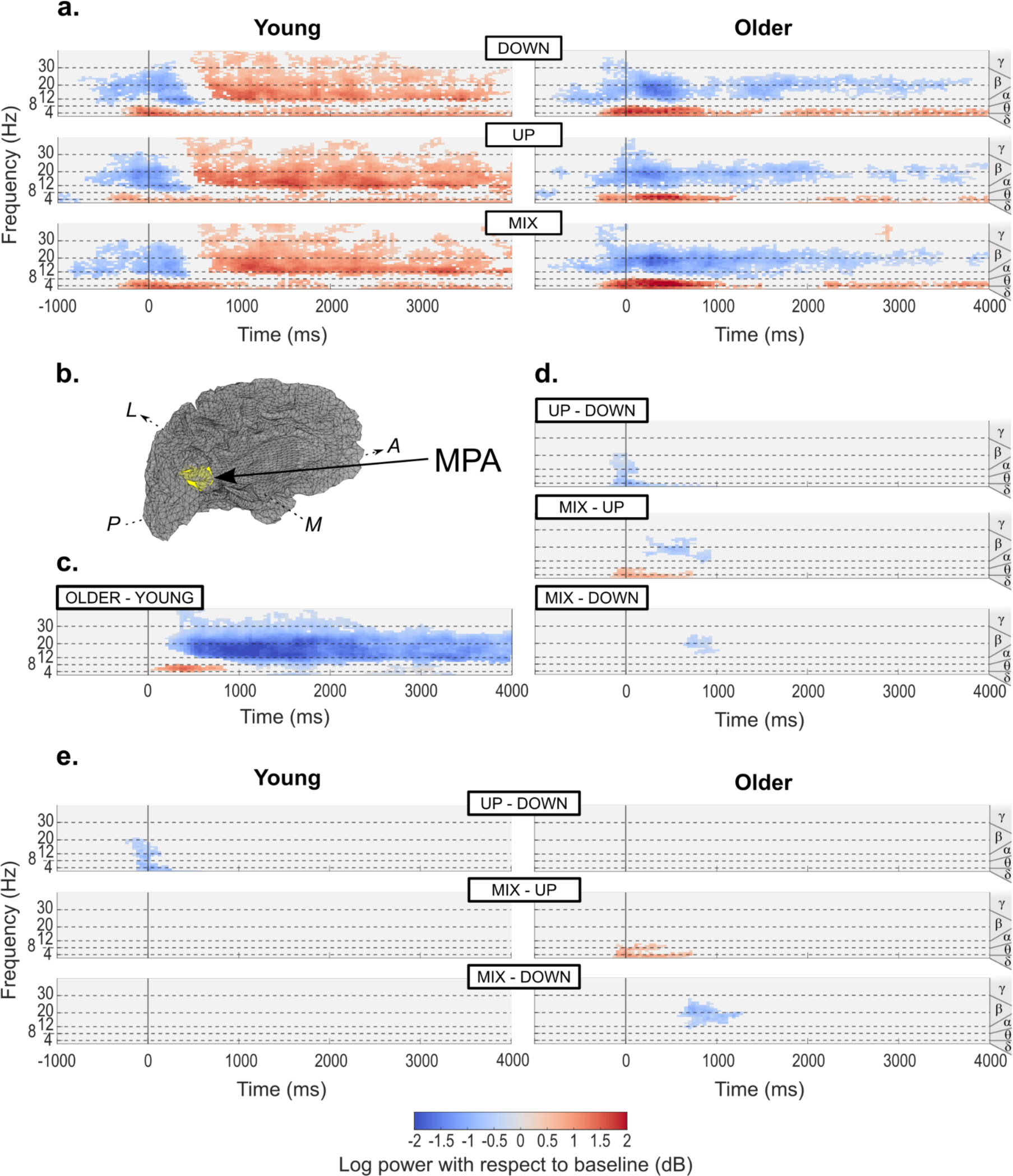
ERSP activity time-locked to the event of arriving at an intersection in the test trials, reconstructed in the MPA. For further details, see Fig. 4 legend. **a**. Average activity per age group and condition baselined with the 2 seconds period prior to arrival at the intersection. We only display activity statistically different from surrogate baseline distribution (*p* < 0.05). **b**. Illustration of the localization of the left MPA overlaid on left hemisphere source space (midgray surface) in one young participant. A: Anterior; P: Posterior; M: Medial; L: Lateral. **c**. Differences between age groups, irrespective of conditions. We only display statistically significant differences (*p* < 0.05). **d**. Differences between conditions, irrespective of age groups. We only display statistically significant differences (*p* < 0.0083, Bonferroni-corrected for 3 two-sided comparisons). **e**. Differences between conditions within each age group. We only display statistically significant differences (*p* < 0.0041, Bonferroni-corrected for 6 two-sided comparisons).

In all ROIs, we also observed significant differences in the lower frequency band between the conditions irrespective of the age group. In the OPA only, the theta synchronization started earlier, 500 ms prior to arriving at the intersection, in the UP condition than in the DOWN condition (Fig. 4d). We also found a significant decrease of delta/theta synchronization upon reaching the intersection in the UP condition with respect to DOWN and MIX conditions in all ROIs (Figs. 4d, 5d, 6d). This effect lasted until +750ms in the PPA (Fig. 5d) and MPA (Fig. 6d),but stopped at around +250 ms in the OPA (Fig. 4d). When analyzing differences between the conditions within each age group, we revealed a significantly lower synchronization in the UP condition than in the DOWN condition between -100 ms and +250 ms in the OPA (Fig. 4e) and MPA of young participants (Fig. 6e). Finally, we noted a significantly stronger synchronization in the MIX condition than in the UP condition up until +750 ms in the MPA of older adults only (Fig. 6e).

Commonly to both age groups and all ROIs (Figs. 4a, 5a, 6a), we found a strong desynchronization spanning the beta band between -500 ms and +500 ms around the event of arriving at the intersection. Notably, after +200 ms this desynchronization was significantly more pronounced in older adults than in young adults for all ROIs (Figs. 4c, 5c, 6c). While the beta desynchronization continued after +500 ms in the older group, we observed an inversion leading to a strong power synchronization in the young group (Figs. 4a, 5a, 6a). This age-related divergence between activity patterns remained consistent until the end of the 4-second window (Figs. 4c, 5c, 6c). In the three ROIs of young participants, and particularly in the OPA, a significant low gamma band (30-40 Hz) synchronization accompanied the age-specific beta band synchronization (Figs. 4a, 5a, 6a). This activity remained significantly more synchronized in young adults than in older adults between +500 ms and +2500 ms (Figs. 4c, 5c, 6c). Differences between conditions were more specific to the ROI in this range of frequencies than in lower frequencies. In the OPA (Fig. 4d) and MPA (Fig. 6d), we found a significantly greater desynchronization in the low beta band (12-20 Hz) for the UP condition than for the DOWN condition around the synchronizing event. Breaking down the analysis within age groups, we reported that this significant difference was mostly attributable to young participants (Figs. 4e, 6e). In the OPA just before +500 ms, we also noted a significantly greater beta band desynchronization in the DOWN and MIX conditions compared to the UP condition (Fig. 4d). In the MPA between +500 ms and +1000 ms, there was a significantly more pronounced beta band desynchronization in the MIX condition than in the UP and DOWN conditions (Fig. 6d). Finally, we found a more sustained beta band desynchronization in the MIX condition than in the DOWN condition between +500 ms and +1000 ms, in the OPA, PPA and MPA of older participants (Figs. 4e, 5e, 6e).

##### Mean ERSP activity per frequency band across ROIs

In a follow-up analysis of activity during the test phase, we sought to compare time-resolved spectral activity between ROIs. We grouped the ERSP activity by frequency band: delta/theta (2-8 Hz), alpha (8-12 Hz), beta (12-30 Hz), and low gamma (30-40 Hz). Within each frequency band we averaged the ERSPs over the spectral dimension, yielding a time-course of mean spectral activity for each ROI across conditions. Using an independent model for each age group, we then performed pairwise statistical analysis with paired permutation tests based on the maximum cluster-level statistic using 1,000 permutations. We set the significance level to *p* < 0.00104, Bonferroni-corrected for 24 two-sided comparisons. We displayed results pertaining to the alpha band in Supplementary Fig. 4 due the absence of significant activity observed in this frequency range in the previous analysis.

In the lowest frequencies, we observed a strong increase in power synchronization around the time of arrival at the intersection across all ROIs and in both age groups (Figs. 7a,b). In the OPA and MPA only, we found the peak of synchronization to be significantly higher and sharper in young participants compared to older participants (Figs. 4c, 6c). In the young group, we reported the delta/theta burst of activity to be earlier and more transient in the OPA than in other ROIs (Fig. 7a). The synchronization in this region fell significantly below that found in the PPA and MPA during the first 1.5 seconds of observation (Fig. 7a). In the older group, the peak width and latency was similar across ROIs but the amplitude of the power synchronization was significantly higher in the MPA than in the OPA and PPA (Fig. 7b). Moreover, we found that in older participants there was a sustained delta/theta synchronization until the end of the observation window that was significantly more important in the PPA and MPA than in the OPA (Fig. 7b).

**Figure 7.**
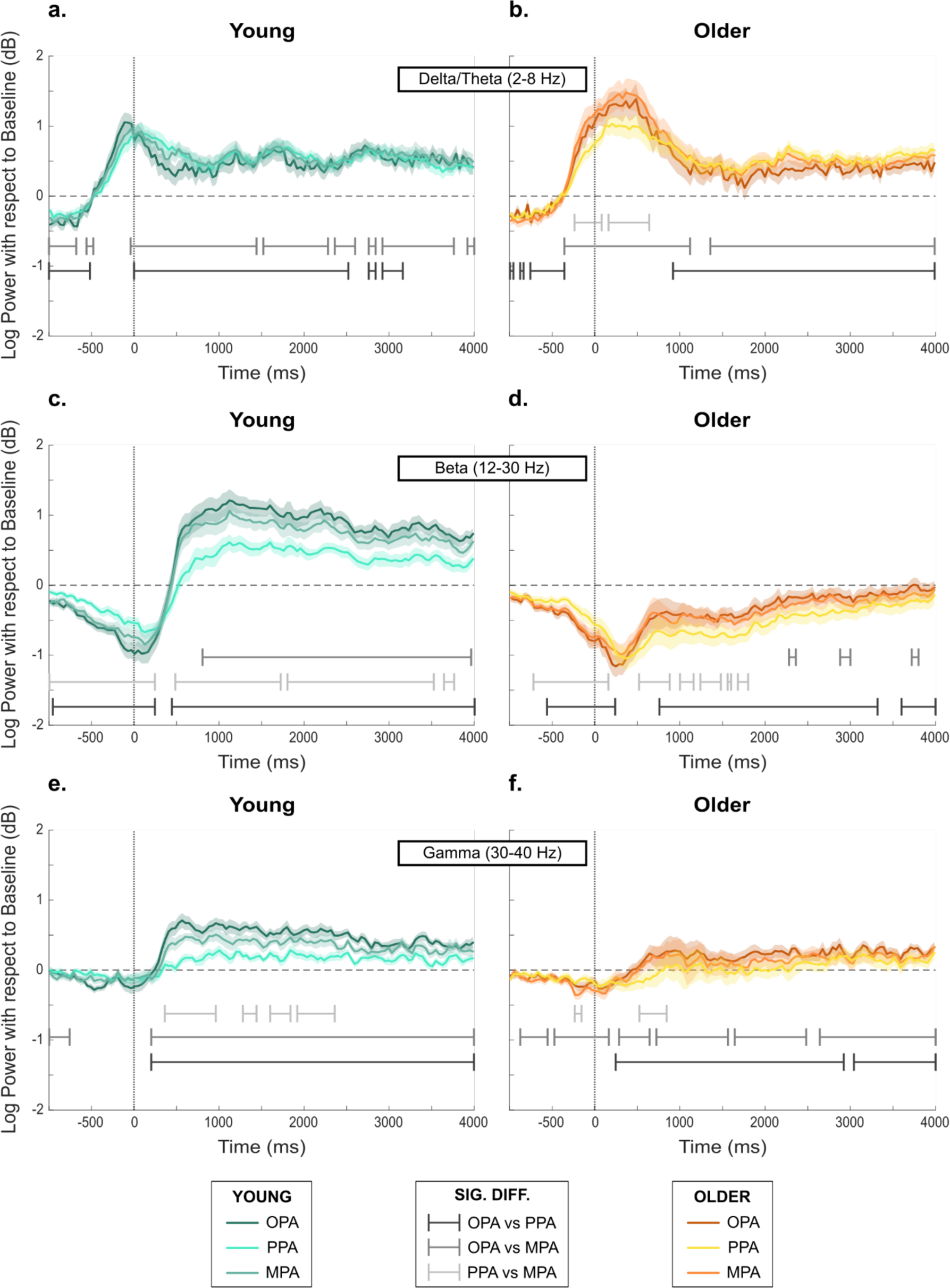
Mean ERSP activity per ROI and frequency band in the test trials across age groups. Bold lines represent the mean ERSP activity per ROI, averaged on the spectral dimension and across all test trials in the corresponding age group. Shaded areas represent the standard error of the mean computed with average subject traces in each age group (n = 21 for each age group). The signal is time-locked to the event of arriving at an intersection (t = 0 ms). We investigated all statistical differences with paired permutation tests based on the maximum cluster-level statistic using 1,000 permutations. To evaluate the statistical significance at the sample level (mean spectral power at a time point) for a given permutation, we used linear mixed-effects modeling. We examined pairwise differences between ROIs for each age group (6 comparisons). Significant differences reflect clusters with a Monte Carlo *p*-value below 0.00104 (Bonferroni-corrected for 24 two-sided comparisons) **a.** ERSP activity averaged over the delta/theta band (2-8 Hz) in the young group. **b.** ERSP activity averaged over the delta/theta band in the older group. **c.** ERSP activity averaged over the beta band (12-30 Hz) in the young group. **d.** ERSP activity averaged over the beta band in the older group. **e.** ERSP activity averaged over the low gamma band (30-40 Hz) in the young group. **f.** ERSP activity averaged over the low gamma band in the older group.

In the beta band, we observed a strong desynchronization starting 1s prior to the arrival at the intersection and peaking at around +250 ms in both age groups (Figs. 7c,d). The peak of desynchronization was slightly delayed but deeper in older adults than in young adults across all ROIs (Figs. 4c, 5c, 6c). However, this early beta desynchronization was less pronounced in the PPA than in the OPA and MPA in both age groups (Figs. 7c,d). After +750 ms, in older participants the beta desynchronization slowly went back to baseline, particularly in the OPA and MPA (Fig. 7d). In young participants, we noted an inversion around +500 ms, leading to a strong beta synchronization in all ROIs (Fig. 7c). Although the activity pattern after +500 ms was more similar between the OPA and the MPA, we reported significant differences between all pairs of ROIs until the end of the observation window (Fig. 7c). Overall, the beta synchronization was strongest in the OPA and weakest in the PPA, with an intermediate activity in the MPA (Fig. 7c).

In the low gamma band, we previously reported a significant synchronization shortly after the arrival at the intersection that was specific to the OPA and MPA of young participants (Figs. 4a, 5a, 6a). The low gamma band activity was significantly more synchronized in the OPA than in the MPA and PPA for the whole course of the observation period in young participants (Fig. 7e). Gamma synchronization in the MPA was significantly stronger than in the PPA during the [+500; +2500 ms] period (Fig. 7e). We revealed a similar trend in the older participant group, albeit on a smaller scale (Fig. 7f). Gamma synchronization was strongest in the OPA and weakest in the PPA, with the MPA in between (Figs. 7e,f).

## Discussion

This work brings together an unprecedented set of tools to shed light on how the vertical position of objects ties in with navigational behavior in young adulthood and healthy aging. We found that young adults’ equivalent performance across conditions echoed with an adaptation of their gaze patterns to the position of relevant cues in the environment. On the contrary, older adults displayed impaired navigational behavior in all conditions and a systematic oculomotor bias for objects in the lower AOI. Source-reconstructed EEG activity revealed extensive age-related differences in the OPA, PPA and MPA. Moreover, we found that the scene-selective regions of young and older adults displayed theta and beta band specificities for environments that contained relevant cues only in the upper AOI.

### Healthy aging is associated with a downward bias in gaze dynamics

We first revealed similar behavioral performance in young participants across conditions. We argue that young adults’ navigational behavior is malleable to variations in the vertical location of relevant objects. This finding is coherent with recorded gaze metrics. Indeed, the distribution of first fixated AOIs highlighted that young adults divided their initial fixations equivalently between lower, central and upper portions of the screen throughout the encoding phase. Along the same lines, young participants spent less time looking at the upper AOI in the DOWN condition than in the other two conditions during the test trials, conveying the idea that their gaze patterns adapted to the condition. These results are in accordance with research showing that the cognitive demands of the task at hand drive the oculomotor patterns in young adulthood (Jovancevic-Misic and Hayhoe, 2009; de Condappa and Wiener, 2016; Grzeschik *et al*., 2019). For example, young adults spend more time gazing at landmarks located at decision points than at non-relevant objects (Hamid, Stankiewicz and Hayhoe, 2010). In line with numerous accounts of navigational deficits in healthy aging, we found that older adults made significantly more errors on the task than their younger counterparts (Moffat, 2009; Lithfous, Dufour and Després, 2013; Lester *et al*., 2017). We reported that strategy correlated with behavior and that few older participants used a map-based strategy to get their bearings in the square-shaped virtual environment. They also displayed better wayfinding skills on repeated routes than on new ones suggesting a sequence-based route learning strategy. A large body of literature has indeed established that difficulties in creating and using cognitive maps for orientation are characteristic of healthy aging (Moffat, Elkins and Resnick, 2006; Wiener, Kmecova and de Condappa, 2012; Allison and Head, 2017; Wiener *et al*., 2020), particularly in environments devoid of geometric information (Bécu *et al*., 2023). The unsuccessful map use in aging may have not been the only contributing factor to such navigational impairments. Indeed, older individuals reported using all 4 objects at intersections, confirming previous research that directing gaze patterns towards the informative part of a scene is frequently impacted in aging (Grzeschik *et al*., 2019; Hilton *et al*., 2020). We hypothesize that older participants experienced visual working memory overload as navigationally irrelevant objects competed with relevant ones (Gazzaley *et al*., 2008; Schmitz, Cheng and De Rosa, 2010). Such behavioral deficits resonate with the observed stereotyped oculomotor behavior. Older participants’ first fixations preferentially targeted the central AOI during the UP condition, neglecting the upper AOI. Moreover, the VMA index revealed that older adults spent more time looking at the lower half of the screen regardless of the condition. In contrast to young participants who modulated their attentional focus according to vertical position of relevant cues in the test phase, older participants displayed a systematic downward gaze bias. Past studies revealed that older people tend to focus on lower portions of a visual scene during locomotion (Uiga *et al*., 2015; Bécu *et al*., 2020). In real-life settings, fixations directed towards the lower field could aid in anticipating stepping patterns, planning trajectories and encoding the geometric layout of the space. Moreover, recent work shed light on an age-related spatial memory deficit that was specific to the upper visual field (Durteste *et al*., 2023). We speculate that the observed vertical shift, apparent both in natural and desktop-based conditions, could represent a default state aimed at minimizing task error in aging.

### A specific neural pattern in the OPA, PPA and MPA during the UP condition

The EEG signatures around the time of arrival at the intersection were equivalent in both age groups. We found strong beta desynchronization along with the transient theta synchronization in all ROIs, in line with a previous study that had participants recalling spatial relationships between objects (Rondina *et al*., 2019). We thus argue that this pattern of cortical activity reflects visual processing of a scene and the objects contained within it. According to the status quo theory, beta band desynchronization in sensory areas is a sign of disruption of the internal state by a strong exogenous stimulation (Engel and Fries, 2010). Moreover, multiple reports demonstrate that transient parietal and medial temporal theta synchronization accompany the processing of landmarks in paradigms assessing spatial cognition (Kober and Neuper, 2011; White *et al*., 2012). Across both age groups, we also observed that the pattern of theta synchronization and beta desynchronization was different in the UP condition than in the DOWN and MIX conditions. This result hints at a specificity of the UP condition; the absence of navigationally relevant objects in the lower half of a scene seems to impact the activity of scene-selective regions. With regards to the fact that all 3 conditions contained an equal quantity of visual information, we emphasize that the vertical position of information useful for navigation modulates cortical signals in scene-selective systems. The vertical retinotopic preferences of the OPA and PPA cannot explain the latter finding as all three ROIs exhibited similar EEG signatures during the UP condition (Silson *et al*., 2015). Nonetheless, early stages of visual processing are tuned to the typical vertical position of objects within the visual field (Kaiser, Moeskops and Cichy, 2018). For example, the upper visual field provides orienting information from distal and large immovable objects whereas the lower visual field frequently contains smaller objects and obstacles (Greene, 2013; Hafed and Chen, 2016; Groen, Silson and Baker, 2017; Saleem, 2020). In our paradigm, we removed distal information by introducing fog which forced participants during the UP condition to orient using proximal objects on balconies. It is therefore conceivable that such condition-associated differences emerged from the atypical scene grammar of intersections in the UP condition. We highlight that this pattern was consistent between age groups, in accordance with a recent report providing evidence for a robust object-scene congruency effect in the OPA and PPA across the lifespan (Rémy *et al*., 2020). Further research should seek to elucidate the mechanisms that confer the UP condition its specificity.

### Age-related differences in oscillatory dynamics reflect discrepancies in attentional processing

Despite these initial similarities, after +500 ms, we reported striking disparities in cortical activity patterns between young and older participants across conditions. The young group displayed a sudden and sustained beta/gamma synchronization which was absent in the older group. This activity has been associated with tasks that require an effort to maintain the current cognitive state (Deiber *et al*., 2007; Engel and Fries, 2010). A previous study has also shown that beta band synchronization over the occipital cortex is a marker of visual attentional processes and that healthy older adults demonstrate a specific reduction of this oscillatory pattern (Gola *et al*., 2013). Similarly, there is converging evidence that gamma oscillations in parieto-occipital regions promote sharper visuospatial attention (Jensen, Kaiser and Lachaux, 2007). We interpret this age-specific beta/gamma synchronization as a sign of top-down modulation, possibly highlighting the reactivation of pre-existing visual representations (Osipova *et al*., 2006). In our opinion, the absence of such a component in the activity of scene-selective regions in older adults might relate to their inability to compare incoming visual information with internal spatial representations. In the older group, instead of the sudden beta synchronization, we found the initial beta desynchronization and delta/theta synchronization to persist until the end of the time window. The intensity of cognitive processing positively correlates with delta power (Harmony, 2013; Güntekin and Başar, 2016). In addition, evidence points towards the amplitude of theta synchronization in temporal regions being associated with impaired, more effortful, spatial encoding (Fellner *et al*., 2016; Rondina II *et al*., 2019). We speculate that the EEG signals in the older group compared with the young group demonstrate prolonged and stronger bottom-up processing evoked by the arrival at the intersection. The sustained synchronized theta power and desynchronized beta power observed in older participants resonate with a recent study putting forward the possibility that this activity underpins the recollection of superfluous contextual details (Strunk *et al*., 2017). In further support of this hypothesis, we noted the greatest beta desynchronization in the condition characterized by the higher number of relevant objects for navigation in the older group (*i.e.,* the MIX condition). Healthy aging correlates with an increased reliance on non-pertinent information, a process known as hyper-binding (Kim, Hasher and Zacks, 2007; Campbell, Hasher and Thomas, 2010). Hyper-binding originates from a failure of the inhibition mechanisms that regulate attention. Taken together, these results shed light on age-related modifications of attentional mechanisms that may have impeded efficient visual processing and ultimately hindered the formation of adequate spatial representations in older participants.

### Three distinct scene-selective systems

The present study aligns with extensive research highlighting functional dissociations between the OPA, PPA and MPA (Epstein and Baker, 2019; Dilks, Kamps and Persichetti, 2022). We did not find any evidence for age-related modulations of lower frequencies in the PPA. The PPA processes scene content and layout as the natural statistics of a scene explain a large proportion of its variance (Bonner and Epstein, 2021; Chaisilprungraung and Park, 2021; Dwivedi, Cichy and Roig, 2021; Aminoff and Durham, 2023). The comparable delta/theta synchronization in the PPA across age groups lends credence to the idea that older adults can adequately extract visual information that is devoid of navigational purpose. In the OPA and MPA, however, we noted a stronger delta/theta synchronization in older adults compared to young adults. The OPA is a major node in the network facilitating immediate visually guided navigation (Dilks, Kamps and Persichetti, 2022). The conclusions from previous fMRI studies converge to suggest that OPA activity increases with age in response to spatial navigation difficulties (Lithfous *et al*., 2018; Ramanoël *et al*., 2020). In a more complex cognitive task requiring the formation of a mental map, older adults demonstrated an elevated recruitment of the OPA and MPA in tandem (Diersch *et al*., 2021). Although we lack converging evidence across brain regions, the BOLD response and theta power have been positively correlated during human navigation in the parahippocampal cortex (Ekstrom *et al*., 2009). In this light, we speculate that the increased theta synchronization in the OPA and MPA of older adults fits with the fMRI literature and confirms a compensatory role for the OPA during spatial navigation in healthy aging. In the younger group, we also noticed that the peak of theta synchronization was earlier and more transient in the OPA than in the other ROIs. This finding is consistent with the claim that the OPA can rapidly extract local scene information in order to guide deliberate gaze exploration during prolonged scene viewing (Malcolm *et al*., 2018; Suzuki *et al*., 2021). Our study thus emphasizes the distinct roles of the PPA and OPA and it provides further confirmation of the OPA’s function in automatic scene parsing for immediate navigational purposes (Harel *et al*., 2022). Further distinguishing the 3 scene-selective regions, we reported the late stage synchronization of higher frequencies in young adults to be ROI-specific. Indeed, the OPA and PPA exhibited the most and least synchronized activity respectively, while the MPA found itself in the middle. We argue that this continuum of synchrony may correlate with the amount of connectivity with other brain areas at intersections, as gamma band oscillations structure communication between regions (Lachaux *et al*., 2005; Fries, 2015). In a recent study, researchers observed that the retrosplenial cortex, but not the parahippocampal region, cooperated actively with the hippocampus and prefrontal cortex during map-based navigation (Qi *et al*., 2022). Despite a paucity of research delving into the connectivity of the OPA, its role in identifying immediately navigable paths is essential to inform navigational decisions (Bonner and Epstein, 2017, 2018; Patai and Spiers, 2017). It thus seems reasonable to speculate that the OPA communicates with an extended cerebral network (Baldassano *et al*., 2016; Ramanoël *et al*., 2019). We stress that shedding light on how neural oscillations in the scene-selective systems mediate the use of spatial representations for wayfinding is a critical next step in the field (Kunz *et al*., 2019).

### Limitations and Perspectives

From a methodological standpoint, we must highlight that although the source reconstruction model provides a fine-tuned localization of scene-selective regions, spatial precision is incomparable to that of an fMRI study. This is particularly true for deep cortical regions as increasing depth correlates with a decrease in synchronization of the highest frequency band. It is conceivable that the lesser synchronization reported in high frequencies for the PPA may stem from its location in deep structures of the brain. We note that this drawback does not affect low frequency bands. A second methodological consideration pertains to the choice of baseline for the EEG analysis. The baseline corresponded to a period devoid of task-relevant visual information, when participants moved passively along the street. We thus compare the intersection observation period to a phase that called for optic flow processing. The latter may have confounded reported differences between ROIs as the OPA represents self-motion information to a greater extent than the MPA and PPA (Kamps, Lall and Dilks, 2016; Sulpizio *et al*., 2020). Optic flow emanates primarily from the lower visual field which may have primed the processing of objects on sidewalks (Saleem, 2020). Furthermore, young adults were quicker than older adults to make the correct navigational decision in the test phase, such that they resumed to passive navigation within the 4-second window more often. We acknowledge that age-related differences in EEG activity may have been influenced in part by this behavioral disparity. A final limitation to this study relates to task design. Indeed, we believe that the difficulty of the paradigm prevented behavioral differences between conditions from emerging in older participants. Interestingly, top performers in the older group made fewer navigational errors in the DOWN and MIX conditions than in the UP condition. We put forward the hypothesis that, with an easier wayfinding task, the vertical shift in gaze behavior would have translated into a greater difficulty to orient in the UP condition in aging. Future studies could design an immersive task that would increase the availability of multisensory cues and as a consequence facilitate performance in older adults (Adamo *et al*., 2012). Finally, a virtual reality or real-world protocol in combination with mobile EEG would confer greater ecological validity while preserving access to the neural correlates of spatial cognition (Delaux *et al*., 2021; Djebbara, Fich and Gramann, 2021; Do, Lin and Gramann, 2021; Liang *et al*., 2021).

### Conclusion

The present study reached two main conclusions. First, it revealed age-related disparities in beta/gamma band synchronization within scene-selective regions concurring with a deficient regulation of attentional mechanisms during navigation in aging. Second, it highlighted the need to consider vertical position as an essential object property for spatial navigation throughout adulthood. Older adults exhibited a systematic bias for downward fixations. We stress that vertical biases in eye movements in healthy aging may have detrimental consequences on wayfinding abilities as relevant cues are not uniformly distributed across the visual field. A gaze bias for the lower portion of a scene may prevent the formation of adequate spatial representations by hindering the capacity of older adults to anchor their position to larger immovable landmarks. Finally, in both age groups, orienting with objects situated in the upper portion of the screen modified neural activity in scene-selective regions. We argue that the OPA, PPA, and MPA have a significant role to play in parsing information across vertical hemifields. Further research is required to confirm this coding property across imaging modalities and to elucidate its behavioral correlates across the lifespan.

## Methods

### Participants

A total of 24 young and 23 older adults from the French SilverSight cohort took part in the experiment (Lagrené *et al*., 2019). All subjects met the cohort inclusion criteria: normal cognitive performance on a battery of neuropsychological tests including the Mini-Mental State Examination (Folstein, Folstein and McHugh, 1975), the 3D mental rotation test (Vandenberg and Kuse, 1978) and the perspective-taking task (Kozhevnikov and Hegarty, 2001), no history of sensory, neurological, or psychiatric disorder, and normal or corrected-to-normal eyesight. We excluded 3 young participants and 1 older participant because their fMRI data were of poor quality and thus not usable for the EEG source reconstruction analysis. One other older participant was removed from analyses as he struggled to stay awake during the spatial navigation task. The final analyzable group comprised 21 young, 7 females and 14 males (29.0 ± 4.27 years-old), and 21 older adults, 11 females and 10 males (75.8 ± 3.79 years-old). Of note, 1 young adult had eye tracking data of poor quality and 3 young adults were missing data for the 3D mental rotation task; they were thus removed from these specific analyses. Each participant provided their written informed consent, and the study was approved by the Ethics Committee “CPP Ile de France V” (ID_RCB 2015-A01094-45, CPP N: 16122).

### Virtual environment

We designed the virtual town with Unity3D v2019.2 (Unity Technologies, Inc. San Francisco, California, USA) for the purpose of this experiment. The virtual environment consisted of a series of four intersections arranged in a square-like layout (Fig. 1c). Streets lined with identical buildings connected the four-way intersections, rendering them indistinguishable from one another. Every intersection comprised four corner balconies and four corner sidewalks, with an object placed at each of these eight positions. Notably, all balcony items and all sidewalk items in any given intersection were strictly identical in order to limit the perceptual and mnemonic load. Therefore, a unique set of two items unambiguously characterized an intersection (Fig. 1a). Participants’ visibility was restricted to the front four objects of an intersection with the addition of gray fog (Fig. 1a).

The specific configuration of objects in the virtual environment defined three separate conditions (UP, DOWN and MIX). The UP, DOWN, and MIX conditions differed by the position of navigationally relevant objects at intersections: on balconies only, on sidewalks only or on both (Fig. 2a). Across intersections of an UP environment, the balcony items were different from each other while the sidewalk items were identical. Across intersections of a DOWN environment, the balcony items were identical while the sidewalk items were different. Finally, across intersections of a MIX environment, both balcony and sidewalk objects differed.

We selected objects included in the virtual environments from 3D Warehouse, a website for pre-made 3D models (http://3dwarehouse.sketchup.com/), and from the Unity Asset Store. Congruency with the virtual town was the main criterion for object selection. To reduce saliency differences between items, we normalized their height to 1 virtual meter and homogenized their color and texture. Specifically, we took special care in attenuating vibrant colors and textures. Before launching the experiment, 5 examiners rated objects according to their recognizability. We discarded and replaced items judged to be poorly recognizable. We created 9 instances of the virtual environment (3 per condition) with this set of objects. We did not reuse items across different instances and we presented the same 9 instances to each participant.

### Virtual navigation task

Participants performed a reorientation paradigm in the previously described virtual town. Their overall task consisted in learning the location of a goal in order to retrieve it from different starting positions throughout the environment. The goal changed position across the various instances of the environment (see *Virtual Environment*) but was always situated in the continuity of one of the four streets making up the square-like layout. A single goal position and six possible starting zones characterized an environment. Six different trajectories could therefore lead to the goal: two that passed through one intersection, two that passed through two intersections, and two that passed through three intersections (Fig. 1c).

A single condition comprised an encoding phase and a test phase. During the encoding phase, subjects navigated passively towards the goal with a forward speed of 5 meters per second and a turning speed of approximately 67° per second. They watched first-person videos of a single-intersection trajectory, a two-intersection trajectory, and a three-intersection trajectory through the environment, in a randomized order. These videos ensured that participants passed through every intersection at least once. The videos marked a 4-second pause at the starting position and at the level of each intersection for participants to encode the objects (Fig. 1d). To succeed in the task, participants had to learn the position of the goal with respect to the objects present at each intersection. Once the three routes had been shown, participants were tested on their reorientation ability. During the test phase, subjects navigated passively through the six possible routes. The video paused upon reaching an intersection and subjects decided whether they needed to go straight, left, or right using the keypad. A correct answer led to navigation resuming, while a wrong decision triggered a sound that notified the participant to retry. Subjects could not move along the route until the correct direction had been chosen. Upon finding the goal, a black screen appeared, and the next video started after a 2-second delay. The first three videos shown during the test phase were always the same as those presented during the encoding phase. The last three test videos corresponded to the remaining three possible trajectories, unexplored during the encoding phase. Such a design served to reduce the difficulty of the task by reinforcing the learning of specific routes.

### Procedure

While we placed the conductive gel, subjects read the paper instructions and watched a demonstration of the paradigm. The demonstration video showed the entire course of a condition, with the encoding and test phases. During the simulated test phase, correct and wrong decisions were made in order for participants to assimilate the two types of feedback. Notably, the virtual environment used for the demonstration was strictly identical to the one used for the actual task except for the objects at intersections which were replaced by colored cubes. We also presented participants with a map of the global layout of the environment to facilitate learning, particularly in older adults (Allison and Head, 2017). Following the EEG preparation and paradigm familiarization phase, the experimental session began with a 5- or 9-point grid eye tracking calibration depending on the subject’s fixation stability. The gaze patterns of the dominant eye were sampled at 500 Hz throughout the experiment.

Subjects performed a total of 9 different conditions as the experiment comprised 3 blocks, each divided into UP, DOWN, and MIX conditions. A block always ended with the MIX condition but could start with an UP or a DOWN condition. Such a configuration mitigated the possibility that subjects starting with the MIX condition would apply the same strategy to perform the UP and DOWN conditions. We counterbalanced the presentation order of the UP and DOWN conditions across blocks and subjects to avoid a potential bias linked to the starting condition. Note that eye tracking calibration was repeated at the beginning of each block, before launching the navigational task, to limit the amount of drift and allow participants to remove their head from the chin rest.

Once the three conditions within a block ended, subjects performed a recognition task. The latter acted as an evaluation of the level of item encoding. Participants saw six objects that they had encountered in the previous virtual environments, two from each condition, and six new objects (*i.e.*, distractors). We presented one item at a time on a black screen, and we randomized their order of presentation. Subjects responded with the keypad whether they believed to have seen the object in one of the three previous environments or not. A pause followed the recognition paradigm, and we adapted its duration to each participant’s individual needs. The experimental session ended with a questionnaire probing subjects’ strategy use and object selection during the reorientation task (Supplementary Fig. 5).

### Apparatus and setting

The virtual navigation task was displayed on a 23-inch Dell monitor with a 1600 x 900-pixel resolution, subtending 49° in width and 30° of visual angle in height. Subjects’ head was positioned 57 cm from the screen on a head-mounted monocular eye tracker (EyeLink 1000 Tower Mount, SR Research Ltd., Canada). The center of the screen was adjusted to be at participant eye-level. Answers were recorded via a numeric keypad (KKmoon) placed on the table in front of participants and adjusted individually for optimal visibility. Subjects wearing progressive lenses removed them for the duration of the task as they could have biased the use of visual information within the upper and lower visual fields. The experimental session took place in a dark room devoid of light, except from that coming from the monitor.

The EEG recording system comprised 127 active wet electrodes mounted on an elastic cap with an equidistant arrangement (waveguard original - ANT Neuro, Hengelo, The Netherlands). An electrode located closest to the standard Cz position (10-20 international system) provided the reference for all other electrodes and an additional one on the left earlobe served as electrical ground reference. After placing the cap on the participant’s head, we performed a head scan to collect the position of individual electrodes using a depth perception camera (RealSense D435 - Intel, Santa Clara, CA, USA) in combination with the RecFusion software (v2.3.0, ImFusion GmbH, München, Germany). We placed visual markers on the location of 3 fiducial landmarks of the skull (nasion and helix/tragus intersection of left and right ears) for later coregistration of the scan with MRI data. Prior to recording for the experimental session, we lowered the impedance of the electrodes with the scalp below 20kΩ. We acquired raw EEG data with eego mylab software (v1.9.1, ANT Neuro) at a 1kHz sampling rate, continuously within each block. To simultaneously record all streams of data (eye tracking, EEG and protocol events), we used the Lab Streaming Layer protocol (Kothe, 2014) and the LabRecorder software (v1.13.0) to collect them into a single XDF file per block.

### Data analysis

#### Behavior

Data were extracted from the XDF files of each participant using the `pyxdf` library in Python 3.8. Analyses were conducted using R version 4.0.3 in RStudio version 2022.12.0+353 (R Core Team, 2020; RStudio Team, 2022). To assess participants’ navigational performance, we computed the number of errors per intersection during the navigational task. This number could vary from 0 to 2, 2 being the total number of wrong directions at a single intersection. We also reported the number of errors made during the recognition task. Using the answers from the post-experiment questionnaire, we grouped participants into two categories: those who used a mental map to orient (from the beginning or at some point during the experiment) and those who never did. As an exploratory analysis, we also classified participants according to the reported number of objects they relied upon for orientation at each intersection: individuals who selected a subset of objects vs. those who relied on all 4 objects. We used one linear mixed model to evaluate the influence of age group, sex, condition, and strategy on performance and a second to test the impact of trajectory type (repeated vs. new) on behavior. We chose the random and fixed effects structures based on the Akaike information criterion (AIC) goodness-of-fit statistic. We carefully inspected the normality of residuals from these models. We also verified using linear regression whether neuropsychological performance as assessed by the Corsi block-tapping, 3D mental rotation, and perspective-taking tasks were associated with performance on the virtual navigation task.

#### Eye-tracking

We conducted all analyses on gaze patterns from the encoding and test phases separately. One young participant had eye tracking data of poor quality and was therefore excluded from the following analyses. We focused solely on the static observation periods at all intersections. The observation period during the test phase depended on the time it took for participants to choose the correct direction. In order to reduce interindividual variability, we restricted our analysis to the first 4 seconds upon arrival at the intersection even when participants took longer to make their decision. We chose a 4-second window in line with the observation phase of the encoding phase. We did not analyze the eye movements during the dynamic parts of videos. The detection algorithm supplied by SR Research classified eye movements into fixations, saccades, and blinks (EyeLink 1000 User Manual, 2005 – 2009). We further processed the data by removing fixations that were shorter than 80 ms or that fell outside the computer screen. We used Gaussian Mixture Modeling (GMM) to cluster fixation data into specific areas of interest (AOIs) for young and older adults separately. We fitted a GMM with n = 5 components to the encoding and test fixation data, as they had the lowest Bayesian Information Criterion (BIC) value. We subsequently labeled each fixation according to the AOI it belonged to, and we visualized the results. For both encoding and test data, we found an AOI at each of the four object positions and an AOI in the center of the screen (Fig. 1b).

Before examining fixation statistics associated with the AOIs, we discarded fixations that fell too far from the center of their attributed AOI. In other words, we removed fixations with coordinates situated outside the ellipse forming the 99th percentile of the Mahalanobis distance (computed for each AOI with the parameters found by GMM). We merged the left and right AOIs into “upper” and “lower” AOIs as we did not have any hypotheses pertaining to left-right asymmetries. We measured three metrics at each intersection: (1) the proportion of time spent looking at each AOI (i.e., the sum of all fixation durations), (2) the first fixated AOI per intersection, and (3) the vertical meridian asymmetry (VMA) related to total dwell time using the following formula:

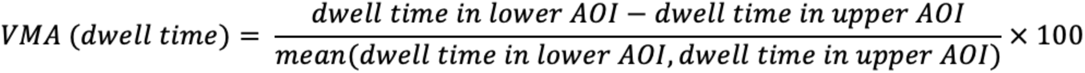

The VMA computes the difference in dwell time between the lower and upper AOIs, it can thus decipher a possible vertical gaze bias. Indices that are close to 0 indicate no asymmetry in dwell time between the upper and lower AOIs. Indices that are below zero reflect an asymmetry in dwell time with more time spent looking at the upper AOI than at the lower AOI. Indices above zero, on the other hand, highlight that more time was spent looking at the lower AOI than at the upper AOI. We ran linear mixed models to assess the impact of age group, sex, and condition on the three eye movement parameters. The latter were summary variables per block and per condition for each participant. We included participant and block as random intercepts in all models, and we compared fixed effect structures, starting with the most complex, using the AIC.

#### EEG

We processed and analyzed the EEG data using Matlab R2019a (Mathworks Inc., Natick, MA, USA) and standalone toolboxes.

##### Source reconstruction modeling

We relied on a fMRI informed source reconstruction model tailored to each participant (Cottereau, Ales and Norcia, 2015). Participants completed fMRI experiments as part of the SilverSight cohort study, we therefore had MRI anatomical scans and relevant fMRI localizer data at our disposal to build such a model.

MRI data was collected using a 3 Tesla Siemens MAGNETOM Skyra whole-body system (Siemens Medical Solutions, Erlangen, Germany) equipped with a 64-channel head coil. Acquisitions took place at the Quinze-Vingts National Ophthalmology Hospital in Paris. For anatomical scans, we used 2 types of sequences: a T1*-weighted MPRAGE 3D sequence (voxel size = 1 x 1 x 1.1 mm, TR/TE/Angle flip = 2300 ms/2.9 ms/900 ms/9°, matrix size = 256 x 240 x 176) and a T2-weighted 3D sequence SPACE CAIPI (voxel size: 1 x 1 x 1.1 mm, TR/TE = 3140 ms/410 ms, matrix size: 256 x 256 x 176). The T2-weighted sequence was available for the young group only. A T2*-weighted SMS-EPI sequence (voxel size = 2.5 x 2.5 x 2.4 mm, TR/TE/Angle flip = 1000 ms/30 ms/90°, matrix size = 100 x 100, SMS = 2, GRAPPA = 2) was acquired for the localizer task. Please refer to Ramanoël *et al*., 2020 for a detailed description of the paradigm.

First, we processed the T1-weighted image of each participant with FreeSurfer (v.7.1.1, Fischl, 2004) using the recon-all command. The latter estimates the interface between gray and white matter and parcellates the brain according to anatomical atlases. We carefully inspected the automatically generated gray and white matter surfaces and we applied manual corrections where necessary. Second, we computed the inner skull, outer skull and outer skin surfaces required for the three-compartment boundary element method (BEM) using the BET tool from the FSL library (Smith, 2002; Jenkinson, Pechaud and Smith, 2005). We selected this tool to improve the automatic estimation with the T2-weighted image. We further checked the automatically generated surfaces and we applied manual corrections when required.

In parallel, we analyzed the localizer task using the SPM12 toolbox implemented in Matlab R2019a (release 7771, Wellcome Department of Imaging Neuroscience, London, UK). We realigned the functional images and co-registered them to the T1-weighted image. We then computed forward and inverse deformation fields from the individual T1 image to the MNI-152 space. Finally, we normalized all images using the forward deformation field, we resampled the functional images to 3 x 3 x 3 mm voxel size (4th degree B-spline interpolation), and we smoothed them using an 8 mm FWHM Gaussian kernel. We fit the localizer data to a single participant general linear model for block design. The model included five categories of interest (scenes, faces, objects, scrambled objects, rest) as regressors, convolved with the SPM canonical HRF. We also added the 6 movement parameters as regressors of no interest in the model and we high-pass filtered each voxel’s time-series (1/128 Hz cut-off). For each participant, we delineated the PPA, OPA, and MPA using the contrast [Scenes > (Faces + Objects)]. We identified significant supra-threshold voxels on individual t-maps that were closest to the reference coordinates for the scene-selective regions (Ramanoël *et al*., 2020) using false discovery rate correction for multiple comparisons (p < 0.05) at peak-level. We regrouped peak coordinates for each young and older participant into Supplementary Table 1.

We selected the best 2D images of the head scan in the RecFusion software to improve the visibility of the electrodes on the 3D model. We subsequently imported the model in the get_chanlocs plugin (v3.00) of EEGLAB (v2021.0, Delorme and Makeig, 2004) to manually point the positions of the 3 fiducials and of each electrode. We then retrieved the 3D position of the electrodes in a coordinate system defined by the fiducials.

We built the forward model matrix using MNE-Python tools (v0.23, Gramfort, 2013). We first defined the midgray surface halfway between white matter and gray matter surfaces. We used a tessellation of this midgray surface with 10242 regularly spaced vertices (hemisphere-wise) as the basis for our source space. Each of these vertices held a current dipole oriented orthogonally to the cortical surface in order to diminish the number of parameters to be estimated in the inverse procedure (Hämäläinen *et al*., 1993; Cottereau, Ales and Norcia, 2015). We individually tessellated inner skull, outer skull and outer skin surfaces into 5120 faces meshes defining the boundaries between the cerebrospinal fluid and the skull, the skull and the scalp and the scalp and the air, respectively. We assigned a standard 0.33 S/m conductivity to brain/cerebrospinal fluid and scalp compartments. The skull thickens and densifies across the lifespan, which modifies its electrical properties. Taking the latter into consideration we chose to assign age-dependent conductivity measures for the skull. We estimated skull conductivity at 0.02145 S/m for young participants (1:15 ratio around 30 years old) and at 0.0099 S/m for older participants (1:33 ratio around 75 years old) (Michel and Brunet, 2019). We coregistered the electrode positions to the MRI head surface using MNE-python coregister function which finds the rigid body transformation that minimizes the distance between fiducials coordinates in both reference frames. Once in the common reference frame, we projected the electrodes onto the outer skin surface. Ultimately, we combined the source space, the electrical boundaries and the electrode locations to characterize the electric field propagation from the cortical sources to the scalp sensors. This step allowed for the computation of the so-called gain matrix.

We solved the inverse problem using the L2-regularized minimum-norm estimation (MNE) (Hämäläinen *et al*., 1993) with the FieldTrip-lite plugin (v20210601, Oostenveld *et al*., 2011) for EEGLAB. Prior to inversion we normalized the contribution of each dipole in the gain matrix to compensate for the bias of the MNE method towards superficial sources (Lin *et al*., 2004; equation 7, parameter gamma = 0.95). As suggested by Cottereau, Ales and Norcia (2015), we imposed local correlation constraints in the source covariance matrix within each ROI to bias the inverse procedure towards coherent activity within these areas. We then proceeded as described in Cottereau, Ales and Norcia (2015). We chose the regularization parameter individually for each participant using a generalized cross-validation approach (Reeves, 1994) implemented in the regtools toolbox for Matlab (Hansen, 2007).

##### EEG processing

For the individual-level EEG analysis, we used a validated pipeline architecture (Delaux *et al*., 2021). We first aggregated all block files from a single participant and processed the data in a continuous fashion. We downsampled the data to 250 Hz and applied a 1.75Hz high-pass filter to suppress slow drifts in EEG data (zero-phase Hamming windowed finite impulse response filter with 1.5 Hz cut-off frequency and 0.5 Hz transition bandwidth, following recommendation by Klug and Gramann (2021). We removed spectral peaks at 50 Hz, corresponding to power line frequency (implemented by the cleanLineNoise function from the PREP pipeline (v0.55.4, Bigdely-Shamlo *et al*., 2015). We identified noisy channels with automated rejection functions, setting parameters values according to the default recommendations. We then reconstructed the removed channels by spherical interpolation of neighboring channels and we applied re-referencing to the common average. In a subsequent time-domain cleaning, we detected and removed artifactual periods with the APP pipeline (da Cruz *et al*., 2018).

On the cleaned dataset, we performed an independent component analysis (ICA) using an adaptive mixture independent component analysis algorithm (AMICA plugin, v1.5.1, Palmer *et al*., 2008). The ICA was preceded by a principal component analysis reduction to the remaining rank of the dataset taking into account the number of channels interpolated. For each independent component (IC), we computed an equivalent current dipole model with the DIPFIT plugin for EEGLAB (v3.7, Oostenveld and Oostendorp, 2002). We used the ICLabel algorithm (version 1.3, Pion-Tonachini, Kreutz-Delgado and Makeig, 2019) with the “default” option to give an automatic class prediction for each IC. Guided by this automatic prediction, we proceeded to a visual check of candidate IC properties (outputted by pop_prop_extended function of EEGLAB) and kept those that contained sufficient evidence for brain-related features. Finally, we back projected the selected ICs in the channel (*i.e.*, electrode) space.

On the cleaned-with-ICA full dataset (*i.e.*, data from all trials, regardless of whether they were labeled as artifactual periods earlier), we ran a final artifact check with the APP pipeline. We used this analysis to automatically flag any epoch containing at least one data point classified as an artifact by both runs of the APP pipeline. These data points contain transient artifacts that were sufficiently strong to be detected prior to ICA but that were not satisfyingly dealt with after removal of non-brain ICs. Before epoching the dataset we applied a bandpass filter between 1.25Hz and 42Hz (zero-phase Hamming windowed finite impulse response filters: high-pass filter with 1 Hz cut-off frequency and 0.5 Hz transition bandwidth and low-pass filter with 47.25 Hz cut-off frequency and 10.5 Hz transition bandwidth). We then epoched the dataset into [-2,+4] second fixed time windows centered around the event of arriving at an intersection. We considered all intersections excluding start positions. We discarded epochs that were previously flagged as well as those that did not contain any fixations. We present the detailed count of epochs included in the EEG analysis in Supplementary Tables 2 and 3.

For the following analyses, we used functions from the FieldTrip-lite plugin (v20210601, Oostenveld *et al*., 2011) for EEGLAB. At the epoch level, we first computed event-related spectral perturbations (ERSPs) for each channel (with a 25 Hz sampling rate in the temporal domain and with a linear scale between 2 and 40 Hz for the spectral domain) using the ft_freqanalysis function. The latter was based on a frequency-wise combination of Morlet wavelets of varying cycle widths (‘superlet’ method: Moca *et al*., 2021). We discarded data from electrodes for which the position was not determined with sufficient confidence on the 3D model. Then we proceeded to solving the inverse problem with the complex Fourier coefficients as inputs using the ft_sourceanalysis function and the MNE method. To summarize activity from the OPA, PPA and MPA, we averaged power estimations over all dipoles included in the ROI and combined its contribution from both hemispheres. Finally, at the epoch level, we divided the activity by its arithmetic mean over the course of the epoch.

##### Group-level analysis

The group-level analysis focused on 4 main variables of interest: (1) age group, (2) condition, (3) the interaction between age and condition and (4) ROI. Since we used a mixed-effects modeling approach on the epoch-based description of the data: there was no within-subject averaging. We performed separate analyses for the encoding and the test phases of the experiment.

We used the period during which participants are transported along the street before the arrival at the intersection (*i.e.,* prestimulus period) as a baseline for cortical activity during scene observation. We defined the baseline activity independently for each variable of interest, frequency and subject and we computed it as the grand average of the prestimulus period from all epochs. We removed baseline activity from each epoch data using a gain (*i.e.,* divisive) model. We then randomly drew samples from the prestimulus period to build surrogate baseline epochs and statistically compare real epochs with baseline-level activity.

We used log-transformed data for statistical analysis as ERSP sample distribution has a better accordance with Gaussian distribution in that space (Grandchamp and Delorme, 2011). To investigate the specificity of ERSPs to the variables of interest within each ROI (age, condition, and their interaction), we performed a statistical analysis using a non-parametric unpaired permutation test based on the maximum cluster-level statistic (Maris and Oostenveld, 2007) with 1,000 permutations. We used linear mixed-effects modeling to evaluate the statistical significance at the “pixel” level (spectral power at a given time-frequency pair) for a given permutation. In the encoding model, we considered participants and instances of the virtual environment as random intercepts and age, condition, sex, block, VMA and the interaction between age and condition as fixed effects (see Eq. 1). In the test model, we kept the same random effects structure and we included trial type (repeated vs. new), number of errors and their respective interaction with age as additional fixed effects (see Eq.2).

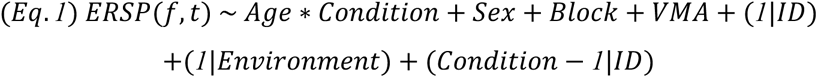

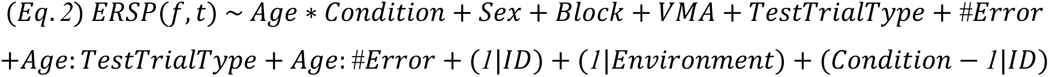

We fit a model for each pixel and we evaluated the contribution of the variable of interest to the model by computing its associated F-statistic. For variables with more than 2 levels, we also extracted pairwise F-statistics. We selected pixels with an F-value above the 95th quantile of the cumulative F-distribution and we clustered them by neighborhood. The cluster-level F-value was the cumulated F-value from all pixels included in the cluster. We then formed the distribution of observed maximum clustered F-values across permutations to compute the Monte Carlo *p*-value for the original repartition. We performed a paired permutation test to assess the statistical significance of ERSP activity for each level of the variables of interest with respect to its corresponding surrogate baseline. Here, we considered participants as random intercepts and original vs. surrogate epochs as a fixed effect (see Eq.3). We set the initial significance level to *p* < 0.05, and Bonferroni-corrected for variables with more than 2 levels.

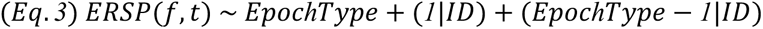

In a follow-up analysis, we sought to investigate the time-resolved spectral activity across ROIs and age groups for all frequency bands: delta/theta (2-8 Hz), alpha (8-12 Hz), beta (12-30 Hz) and gamma (30-40 Hz). We averaged the ERSPs over the spectral dimension at the epoch level, yielding a time-course of mean spectral activity for each frequency band, ROI and age group. To study the influence of ROI in each age group on this signal, we performed a non-parametric paired permutation test based on the maximum cluster-level statistic (Maris and Oostenveld, 2007) with 1,000 permutations. To evaluate the statistical significance at the sample level for a given permutation (mean spectral power at a given time), we used linear mixed-effects modeling for each age group independently. We considered participants and instances of the virtual environment as random intercepts and ROI, condition, sex, block, VMA, trial type, number of errors and the interaction between ROI and condition as fixed effects (see Eq.4).

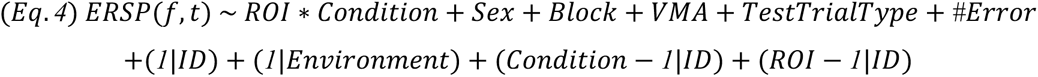

We fit a model for each sample, we estimated the F-statistic for ROI and we extracted pairwise F-statistics. We selected the samples with an F-value above the 95th quantile of the cumulative F-distribution and clustered them by neighborhood. The cluster-level F-value was the cumulated F-value of all samples included in the cluster. Using the distribution of observed maximum clustered F-values across permutations, we found the Monte Carlo p-value for the original repartition. We set the significance level to *p* < 0.00104, Bonferroni-corrected for 24 (2 models x 4 frequency bands x 3 pairwise comparisons) two-sided comparisons.

## Data availability

The datasets generated and analyzed during the current study are available via the OSF repository: https://osf.io/fr5tv/?view_only=413c295de14c477e83cdf1f30e177110

## Code availability

Custom codes used to run the behavioral and eye tracking analyses and to plot the results are available at: https://osf.io/fr5tv/?view_only=413c295de14c477e83cdf1f30e177110. We wrote the codes using R version 4.0.3 in RStudio version 1.4.1103 (R Core Team, 2020; RStudio Team, 2021). All custom software for the EEG analysis can be found at https://osf.io/fr5tv/?view_only=413c295de14c477e83cdf1f30e177110. We wrote the EEG analysis pipeline in Matlab R2019a (Mathworks Inc., Natick, MA, USA).

## Acknowledgements

We would first like to express our most sincere gratitude to all the participants involved in this experiment. We thank Fabienne Tzvetkov-Ricard and Sonia Combariza from the Aging in Vision and Action laboratory at the Vision Institute for their help in enrolling participants. We also wish to thank Bilel Benziane for his insightful suggestions on task design in Unity and Arnaud Kasteler for his help in collecting the data. This research was supported by the French National Research Agency (ANR-18-CHIN-0002), the LabEx LIFESENSE (ANR-10-LABX-65), the IHU FOReSIGHT (ANR-18-IAHU-01), the Fondation pour la Recherche sur Alzheimer (FRA) and the Fondation pour la Recherche Médicale (FRM - FDT202204014752).

## Author contributions

M.D., A.D., Ai.A., S.R. and An.A. designed the experiment. M.D., A.D., Ai.A. and S.R. collected the data. M.D., A.D., B.C. and D.S. analyzed the data. M.D. and A.D. wrote the original manuscript. M.D., A.D. Ai.A., B.C., S.R, D.S. and An.A. revised the manuscript.

