## Supplementary Information for "Age-related disparities in oscillatory dynamics within scene-selective regions during spatial navigation"

### The vertical position of navigational cues drives scene-selective oscillatory dynamics throughout adulthood

### Supplementary Information

#### Supplementary Results

##### ERSP activity in the ROIs

We conducted an ERSP analysis time-locked to the event of arriving at an intersection (t = 0 ms) and grouped by principal frequency bands (delta: <4Hz, theta: 4-8Hz, alpha: 8-12Hz, beta: 12-30Hz, gamma: >30Hz). To summarize bilateral ROI activity, we averaged the source-reconstructed power spectrum from both hemispheres. We reported average ERSPs baselined with the activity preceding the arrival at the level of an intersection. This baseline corresponds to participants passively navigating through a street that is devoid of any relevant spatial information. We computed ERSP statistical differences with permutation tests based on the maximum cluster-level statistic (Maris and Oostenveld, 2007) using 1,000 permutations. We set the initial significance level to $p < 0.05$, and Bonferroni-corrected for variables with more than 2 levels.

###### Encoding trials

The investigated ERSP activity during the encoding trials relates to the perception of the intersection and the objects contained within it. We provided a fixed 4-second static observation period at each intersection to give participants enough time to scan the whole scene and encode the spatial relationships between intersections (Fig. 1d).

We found a strong synchronization in the delta/theta frequency band (2-8 Hz) across all ROIs in both age groups, starting 250 ms before event onset and sustained until +2000 ms (Supplementary Figs. 1a, 2a, 3a). In the MPA, this activity was significantly more pronounced for older adults compared to young adults. Indeed, we reported an increase in theta synchronization (4-8Hz) around the first second of observation, between +400 ms and +1400 ms (Supplementary Fig. 3c). We observed no significant differences between conditions for this range of frequencies in any of the ROIs, neither across groups nor within groups (Supplementary Figs. 1d,e, 2d,e, 3d,e).

Commonly to both age groups and all ROIs (Supplementary Figs. 1a, 2a, 3a), we found a desynchronization in the beta band (12-30 Hz) between -500 ms and +500 ms, around the event of arriving at the intersection. In the PPA, this desynchronization started slightly later, at around -250 ms, and it was mostly restricted to the lower beta band (12-20 Hz, Supplementary Fig. 2a). The beta desynchronization persisted after +500 ms in the older group for all ROIs. In the young group, however, we observed an inversion leading to a power synchronization in the OPA and MPA and an interruption of the desynchronization in the PPA (Supplementary Figs. 1a, 2a, 3a). Notably, this significant age-related difference in the beta band was sustained until the end of the 4-second window in the three ROIs (Supplementary Figs. 1c, 2c, 3c). Although in the PPA it remained mostly localized to the lower beta band (Supplementary Fig. 2c), in the OPA and MPA it extended to the low gamma band (30-40 Hz; Supplementary Figs. 1c, 3c). The low gamma band activity remained significantly more synchronized in young participants than in older participants from event onset until +3500 ms in the OPA (Supplementary Fig. 1c), and from event onset until +2000 ms in MPA (Supplementary Fig. 3c). We observed no significant differences between conditions, whether across groups or within groups, for this range of frequencies in any of the ROIs (Supplementary Figs. 1d,e, 2d,e, 3d,e).

#### Test trials

##### *Mean ERSP activity per frequency band across ROIs*

In a follow-up analysis of activity during the test phase, we sought to compare time-resolved spectral activity between ROIs. We grouped the ERSP activity by frequency band: delta/theta (2-8 Hz), alpha (8-12 Hz), beta (12-30 Hz), and low gamma (30-40 Hz). Within each frequency band we averaged the ERSPs over the spectral dimension, yielding a time-course of mean spectral activity for each ROI across conditions. We then performed pairwise statistical analysis with permutation tests based on the maximum cluster-level statistic using 1,000 permutations. We set the significance level to $p < 0.000104$, Bonferroni-corrected for 24 two-sided comparisons. Here, we display results pertaining to the alpha band due the low activity observed in this frequency range (Supplementary Fig. 4).

In the alpha band, we observed a strong desynchronization starting 1s prior to arrival at the intersection and peaking at around +250 ms in the young group (Supplementary Fig. 4a). This early alpha desynchronization differed across SSRs. It was the least pronounced in the PPA,and the most pronounced in the OPA, with the MPA in-between (Supplementary Fig. 4a). Alpha activity in the older group was desynchronized over the entire 4-second window, with a peak around +500 ms (Supplementary Fig. 4b). Desynchronization was overall more pronounced in the OPA than the MPA, particularly in the first 2 seconds after arrival at the intersection (Supplementary Fig. 4b). Similarly to young adults, the PPA was less desynchronized than the OPA and MPA in the 1s period prior to arriving at the intersection (Supplementary Fig. 4b).

#### Supplementary Figures

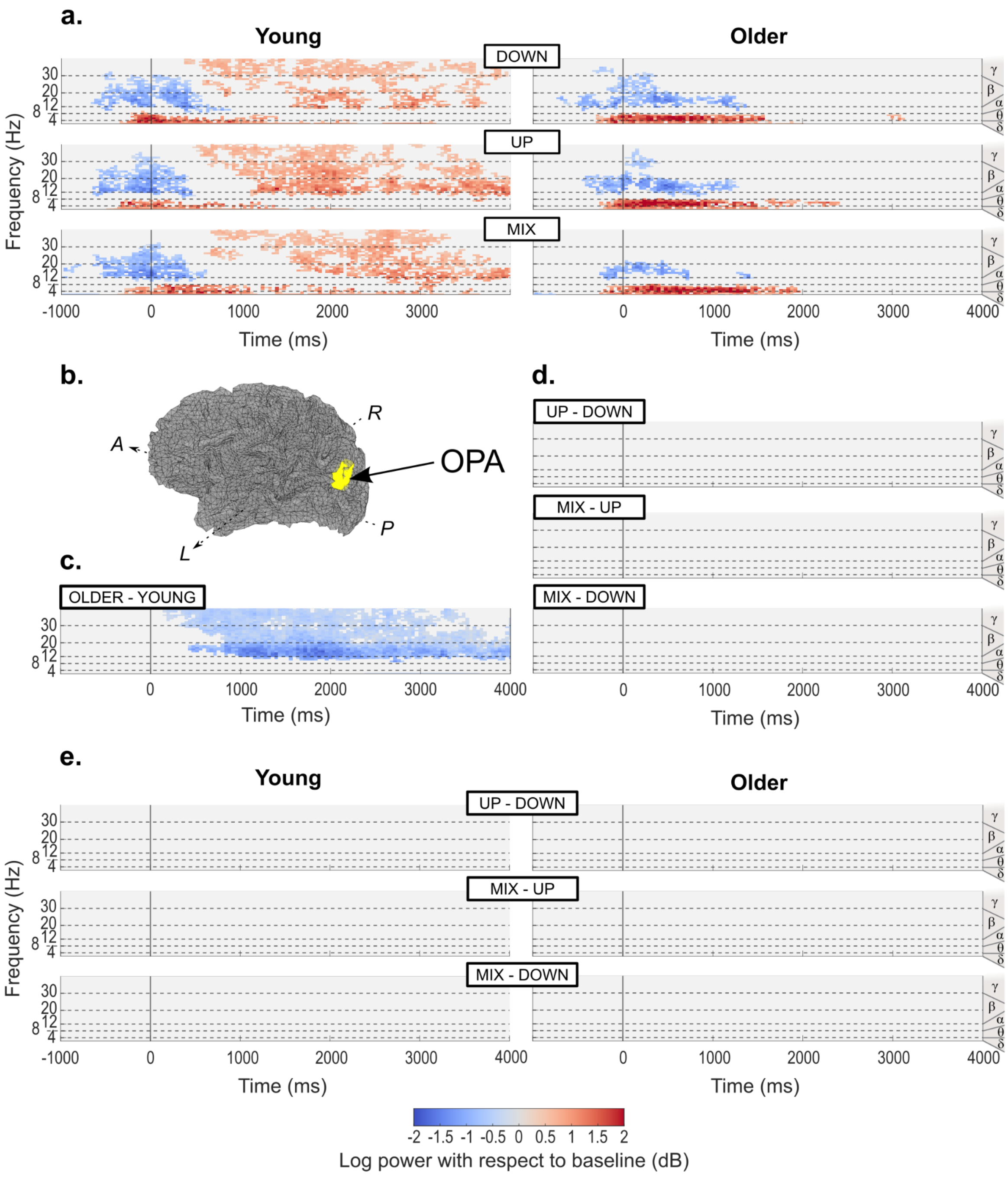

**Supplementary Figure 1**. ERSP activity time-locked to the event of arriving at an intersection in the encoding trials, reconstructed in the OPA. Activity merged from both hemispheric locations after source reconstruction. We examined delta (δ; <4Hz), theta (θ; 4-8Hz), alpha (α; 8-12Hz), beta (β; 12-30Hz) and gamma (γ; >30Hz) frequency bands. We investigated all statistical differences with permutation tests based on the maximum cluster-level statistic using 1,000 permutations. We used linear mixed-effects modeling to evaluate the statistical significance at the "pixel" level (spectral power at a given time-frequency pair) for a given permutation. **a**. Average activity per age group and condition baselined with the 2 seconds period prior to arrival at the intersection. We only display activity statistically different from surrogate baseline distribution ($p$ < 0.05). **b**. Illustration of the localization of the left OPA overlaid on left hemisphere source space (midgray surface) in one young participant. A: Anterior; P: Posterior; M: Medial; L: Lateral. **c**. Differences between age groups, irrespective of conditions. We indicate the direction of the difference above each graph (*e.g.* for the graph entitled "OLDER-YOUNG", we subtracted the average signal for the young adults from the average signal for the older adults, such that a positive difference means more power in the older group than in the young group and a negative difference means less power in the older group than in the young group). We only display statistically significant differences ($p$ < 0.05). **d**. Differences between conditions, irrespective of age groups. We only display statistically significant differences ($p$ < 0.0083, Bonferroni-corrected for 3 two-sided comparisons). **e**. Differences between conditions within each age group. We only display statistically significant differences ($p$ < 0.0041, Bonferroni-corrected for 6 two-sided comparisons).

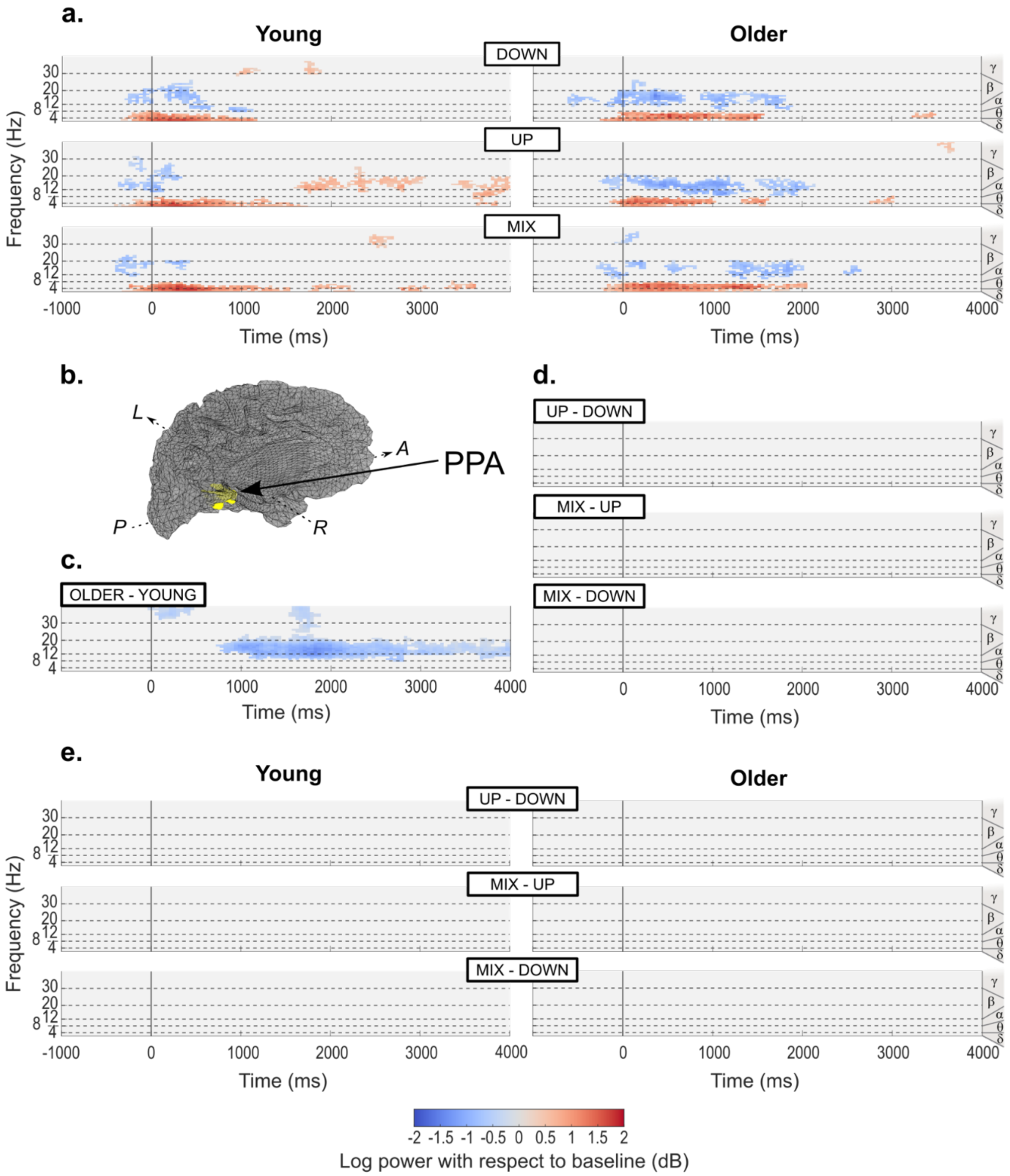

**Supplementary Figure 2**. ERSP activity time-locked to the event of arriving at an intersection in the encoding trials, reconstructed in the PPA. For further details, see Supplementary Fig. 1 legend. **a**. Average activity per age group and condition baselined with the 2 seconds period prior to arrival at the intersection. We only display activity statistically different from surrogate baseline distribution ($p < 0.05$). **b**. Illustration of the localization of the left PPA overlaid on left hemisphere source space (midgray surface) in one young participant. A: Anterior; P: Posterior; M: Medial; L: Lateral. **c**. Differences between age groups, irrespective of conditions. We only display statistically significant differences ($p < 0.05$). **d**. Differences between conditions, irrespective of age groups. We only display statistically significant differences ($p < 0.0083$, Bonferroni-corrected for 3 two-sided comparisons). **e**. Differences between conditions within each age group. We only display statistically significant differences ($p < 0.0041$, Bonferroni-corrected for 6 two-sided comparisons).

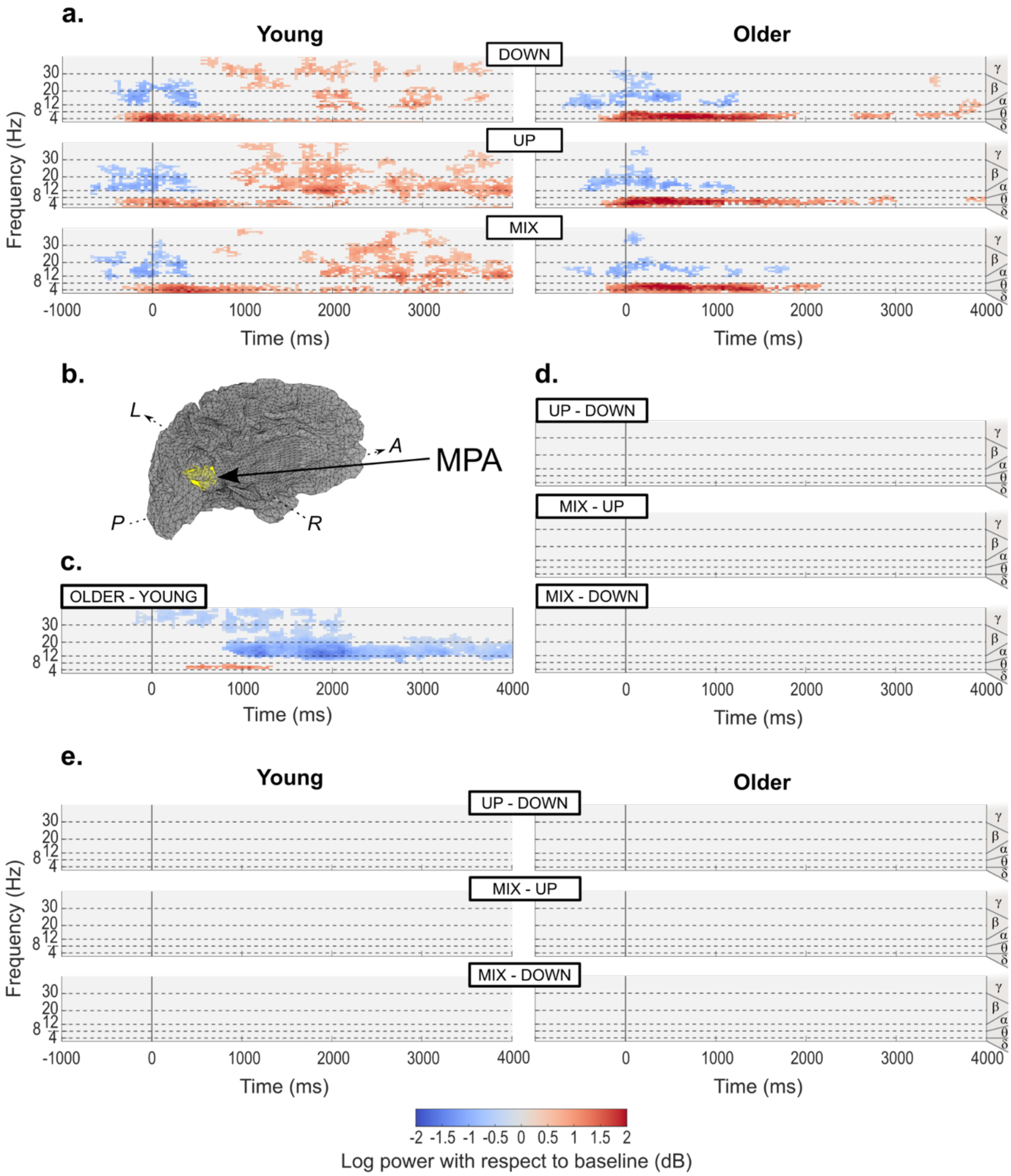

**Supplementary Figure 3**. ERSP activity time-locked to the event of arriving at an intersection in the encoding trials, reconstructed in the MPA. For further details, see Supplementary Fig. 1 legend. **a**. Average activity per age group and condition baselined with the 2 seconds period prior to arrival at the intersection. We only display activity statistically different from surrogate baseline distribution ($p < 0.05$). **b**. Illustration of the localization of the left MPA overlaid on left hemisphere source space (midgray surface) in one young participant. A: Anterior; P: Posterior; M: Medial; L: Lateral. **c**. Differences between age groups, irrespective of conditions. We only display statistically significant differences ($p < 0.05$). **d**. Differences between conditions, irrespective of age groups. We only display statistically significant differences ($p < 0.0083$, Bonferroni-corrected for 3 two-sided comparisons). **e**. Differences between conditions within each age group. We only display statistically significant differences ($p < 0.0041$, Bonferroni-corrected for 6 two-sided comparisons).

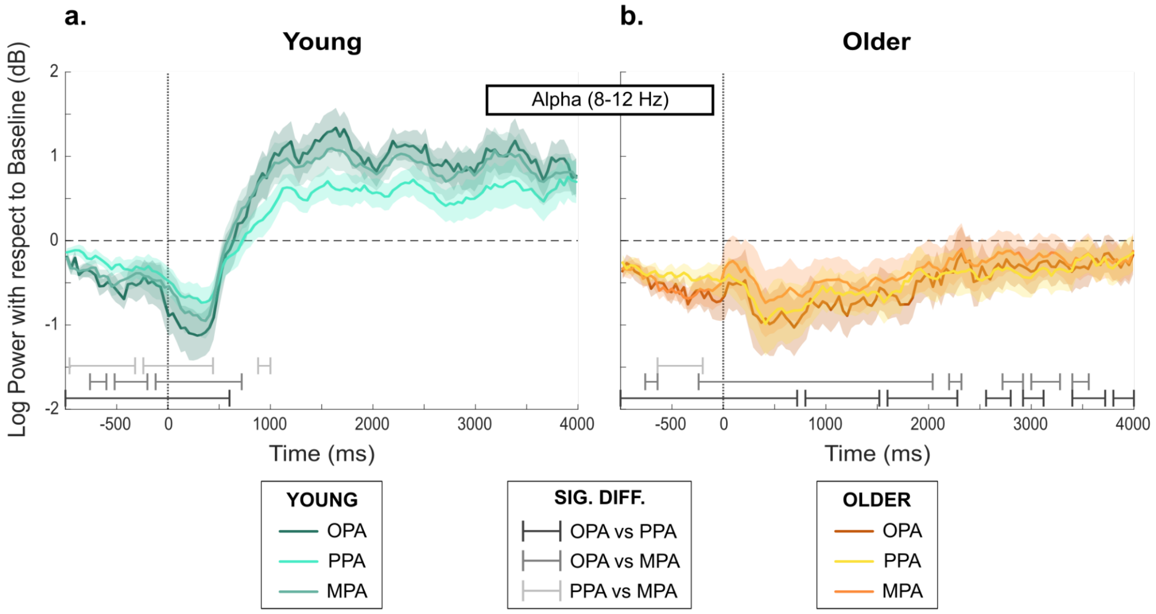

**Supplementary Figure 4**. Mean ERSP activity per ROI in the alpha band (8-12 Hz) in the test trials across age groups. Bold lines represent the mean ERSP activity per ROI, averaged on the spectral dimension and across all test trials in the corresponding age group. Shaded areas represent the standard error of the mean computed with average subject traces in each age group (n = 21 for each age group). The signal is time-locked to the event of arriving at an intersection (t = 0 ms). We investigated all statistical differences with permutation tests based on the maximum cluster-level statistic using 1,000 permutations. To evaluate the statistical significance at the sample level (mean spectral power at a time point) for a given permutation, we used linear mixed-effects modeling. We examined pairwise differences between ROIs for each age group (6 comparisons). Significant differences reflect clusters with a Monte Carlo *p*-value below 0.00104 (Bonferroni-corrected for 24 two-sided comparisons) **a.** ERSP activity in the young group. **b.** ERSP activity in the older group.

#### Questionnaire Post-Expérience

**Différences entre les environnements**

**1 ) Hormis les objets présents aux intersections, avez-vous remarqué d'autres différences entre les environnements ?**

*Cochez toutes les cases qui vous semblent correctes*

- ☐ La longueur des routes
- ☐ La position des objets utiles à l'orientation
- ☐ Les bâtiments de la ville
- ☐ Aucune différence
- ☐ Autre, précisez: ...

**Position des objets utiles à l'orientation**

**2 ) Vous souvenez-vous d'un environnement où les objets utiles à l'orientation étaient...**

*Considérez séparément chaque environnement rencontré et cochez toutes les cases qui vous semblent s'appliquer*

- ☐ Uniquement sur les trottoirs
- ☐ Uniquement sur les balcons
- ☐ Simultanément sur les trottoirs et sur les balcons

**3 ) Comment choisissiez-vous les objets qui vous aidaient à vous repérer à chaque intersection ?**

- ☐ Je choisissais toujours ceux que je préférais
- ☐ Je choisissais toujours ceux sur les balcons
- ☐ Je choisissais toujours ceux sur les trottoirs
- ☐ Je choisissais toujours ceux à gauche
- ☐ Je choisissais toujours ceux à droite
- ☐ Autre, précisez: ...

**Stratégie utilisée**

**4 ) Est-ce qu'une ou plusieurs stratégie(s) suivante(s) se rapproche(nt) de la vôtre ?**

*Cochez toutes les cases qui vous paraissent convenir, en considérant l'expérience dans son ensemble, notamment si vous avez changé de stratégie en cours de route.*

- ☐ (1) J'ai crée une carte vue de haut de l'environnement dans ma tête
- ☐ (2) Je me suis souvenu(e) de l'enchaînement des différentes directions (exemple: "En partant du lion je tourne à droite puis à gauche.")
- ☐ (3) Je me suis souvenu(e) de l'enchaînement des différents objets (exemple: "En partant du lion je croise le buisson puis en tournant à gauche j'arrive au banc.")

**5 ) Si vous avez coché plusieurs cases, diriez-vous plutôt que :**

- ☐ Vous avez utilisé toutes ces stratégies simultanément
- ☐ Vous avez évolué dans votre stratégie, en abandonnant une stratégie pour une autre qui était plus efficace

**Ordre des stratégies utilisées**

**6 ) Vous avez répondu que vos stratégies ont évolué au cours de l'expérience. Sauriez-vous dire comment ?**

*Exemple: "A la moitié de l'expérience, j'ai abandonné la stratégie d'apprendre des enchaînements de tournants pour créer une carte dans ma tête."*

- ☐ Expliquez...

**Supplementary Figure 5**. Post-experiment questionnaire (in French) given to participants as a Google form. The first section ("**Différences entre les environnements**") relates to differences between environments and whether participants perceived them. The second section ("**Position des objets utiles à l'orientation**") relates to the position of objects at intersections and whether participants displayed any preferences. The third section ("**Stratégie utilisée**") relates to the strategy employed by participants to find the goal. Finally, we only displayed the fourth section ("**Ordre des stratégies utilisées**") to participants who had changed strategy during the experiment and asked them to explain this shift.

| Participant | ROI | | | | | |
|---|---|---|---|---|---|---|
| YOUNG | PPA | | OPA | | MPA | |
| | R | L | R | L | R | L |
| **1** | [30, -34,-13] | [-30,-40,-10] | [36,-82,32] | [-36,-85,32] | [15,-52,23] | [-18,-61,20] |
| **2** | [18,-40,-10] | [-27,-43,-10] | [33,-88,23] | [-39,-82,26] | [15,-58,8] | [-18,-61,8] |
| **3** | [27,-40,-13] | [-24,-49,-7] | [36,-85,26] | [-33,-85,14] | [15,-52,11] | [-18,-58,14] |
| **4** | [24,-40,-13] | [-24,-46,-7] | [36,-76,38] | [-27,-85,32] | [18,-52,17] | [-18,-55,14] |
| **5** | [24,-40,-10] | [-24,-46,-7] | [36,-76,35] | [-30,-85,32] | [24,-55,14] | [-24,-55,8] |
| **6** | [27,-52,-7] | [-24,-52,-10] | [36,-79,17] | [-30,-88,20] | [18,-49,11] | [-15,-55,11] |
| **7** | [27,-46,-7] | [-30,-43,-7] | [33,-82,23] | [-33,-82,26] | [18,-52,17] | [-15,-58,20] |
| **8** | [27,-37,-13] | [-24,-43,-13] | [30,-85,32] | [-33,-88,29] | [18,-55,20] | [-18,-61,20] |
| **9** | [24,-46,-7] | [-24,-49,-16] | [36,-76,23] | [-36,-82,20] | [15,-49,14] | [-15,-55,17] |
| **10** | [27,-40,-10] | [-27,-43,-7] | [45,-76,29] | [-39,-79,26] | [21,-55,17] | [-18,-58,20] |
| **11** | [27,-52,-7] | [-24,-49,-10] | [36,-82,32] | [-33,-82,32] | [21,-58,23] | [-15,-61,20] |
| **12** | [27,-46,-7] | [-27,-46,-7] | [33,-79,26] | [-33,-76,20] | [21,-58,14] | [-18,-52,8] |
| **13** | [27,-40,-13] | [-24,-49,-13] | [42,-79,26] | [-39,-82,23] | [21,-52,17] | [-21,-61,14] |
| **14** | [24,-46,-10] | [-27,-49,-10] | [45,-76,17] | [-36,-85,29] | [27,-49,11] | [-24,-61,20] |
| **15** | [33,-58,-7] | [-27,-55,-7] | [39,-73,20] | [-36,-79,23] | [21,-55,14] | [-18,-58,14] |
| **16** | [27,-43,-13] | [-21,-40,-10] | [39,-79,26] | [-39,-76,20] | [21,-55,17] | [-18,-52,8] |

| | | | | | | |
|---|---|---|---|---|---|---|
| **17** | [24,-43,-13] | [-24,-46,-10] | [39,-82,14] | [-39,-82,14] | [15,-52,8] | [-15,-55,8] |
| **18** | [27,-46,-7] | [-24,-49,-7] | [33,-82,26] | [-30,-82,23] | [21,-58,20] | [-18,-55,17] |
| **19** | [27,-43,-10] | [-27,-46,-4] | [39,-79,23] | [-33,-85,23] | [21,-55,14] | [-18,-55,11] |
| **20** | [30,-46,-7] | [-30,-43,-10] | [33,-76,35] | [-33,-79,32] | [21,-52,17] | [-18,-55,17] |
| **21** | [27,-49,-16] | [-27,-46,-13] | [45,-76,17] | [-42,-79,20] | [18,-58,20] | [-18,-55,17] |
| OLDER | **PPA** | | **OPA** | | **MPA** | |
| | R | L | R | L | R | L |
| **22** | [24,-40,-10] | [-27,-43,-10] | [33,-79,32] | [-33,-82,35] | [18,-52,17] | [-18,-61,23] |
| **23** | [30,-43,-10] | [-27,-43,-13] | [48,-79,17] | [-51,-76,11] | [18,-55,11] | [-27,-61,17] |
| **24** | [27,-52,-10] | [-27,-37,-16] | [36,-85,29] | [-30,-88,29] | [21,-55,17] | [-18,-58,20] |
| **25** | [27,-40,-10] | [-24,-37,-13] | [42,-79,23] | [-36,-82,23] | [18,-49,20] | [-12,-52,14] |
| **26** | [24,-40,-10] | [-24,-40,-13] | [36,-76,26] | [-33,-82,26] | [12,-52,11] | [-12,-55,8] |
| **27** | [24,-43,-10] | [-30,-37,-10] | [33,-85,32] | [-39,-82,29] | [18,-55,20] | [-21,-58,23] |
| **28** | [27,-37,-13] | [-27,-40,-10] | [45,-73,20] | [-33,-79,20] | [18,-52,17] | [-15,-61,17] |
| **29** | [30,-49,-4] | [-21,-40,-7] | [36,-85,32] | [-36,-85,29] | [15,-52,17] | [-21,-61,14] |
| **30** | [30,-34,-13] | [27,-43,-10] | [39,-73,17] | [-33,-82,23] | [24,-55,20] | [-21,-61,20] |
| **31** | [30,-55,-10] | [-27,-40,-13] | [33,-76,26] | [-36,-85,26] | [27,-58,20] | [-18,-55,20] |
| **32** | [21,-40,-10] | [-24,-49,-13] | [36,-79,32] | [-33,-85,29] | [18,-55,20] | [-15,-58,20] |

| | | | | | | |
|---|---|---|---|---|---|---|
| **33** | [27,-43,-13] | [-21,-43,-13] | [30,-82,26] | [-21,-88,29] | [18,-58,23] | [-12,-61,14] |
| **34** | [24,-37,-10] | [-21,-37,-13] | [36,-79,26] | [-30,-85,23] | [18,-55,20] | [-15,-70,14] |
| **35** | [27,-40,-10] | [-27,-46,-7] | [36,-79,29] | [-33,-79,23] | [18,-55,20] | [-15,-61,14] |
| **36** | [24,-49,-10] | [-24,-43,-13] | [36,-85,29] | [-39,-85,29] | [12,-52,14] | [21,-58,20] |
| **37** | [27,-49,-13] | [-27,-52,-16] | [33,-82,35] | [-33,-79,35] | [21,-49,14] | [-18,-55,17] |
| **38** | [30,-40,-7] | [-24,-43,-4] | [39,-79,20] | [-36,-79,23] | [15,-49,11] | [-18,-52,11] |
| **39** | [30,-49,-10] | [-27,-43,-10] | [42,-82,23] | [-33,-79,26] | [12,-55,17] | [-15,-61,14] |
| **40** | [33,-34,-13] | [-24,-43,-7] | [39,-73,35] | [-36,-76,23] | [15,-55,11] | [-9,-55,8] |
| **41** | [27,-52,-7] | [-24,-46,-7] | [36,-82,29] | [-30,-88,26] | [21,-52,14] | [-18,-58,8] |
| **42** | [24,-55,-10] | [-27,-43,-7] | [30,-82,23] | [-30,-85,23] | [18,-55,17] | [-18,-55,23] |

**Supplementary Table 1**. Coordinates [X, Y, Z] in the Montreal Neurological Institute (MNI)-152 space of the peak voxel (highest t-value) for the right (R) and left (L) scene-selective regions: PPA, OPA, and MPA for each young and older participant included in the study.

| Num epochs in encoding | UP | MIX | DOWN | Total |
|---|---|---|---|---|
| YOUNG | 344 | 347 | 346 | 1037 |
| OLD | 350 | 343 | 349 | 1042 |
| Total | 694 | 690 | 695 | 2079 |

**Supplementary Table 2.** Number of epochs in the encoding phase used for the ERSP analysis, detailed by age group and condition.

| Num epochs in test | UP | MIX | DOWN | Total |
|---|---|---|---|---|
| YOUNG | 633 | 631 | 632 | 1896 |
| OLD | 663 | 672 | 675 | 2010 |
| Total | 1296 | 1303 | 1307 | 3906 |

**Supplementary Table 3.** Number of epochs in the test phase used for the ERSP analysis, detailed by age group and condition.
